# Invasive alien insects represent a clear but variable threat to biodiversity

**DOI:** 10.1101/2022.06.16.496186

**Authors:** David A. Clarke, Melodie A. McGeoch

**Author notes:** Corresponding author: David A. Clarke.

## Abstract

Invasive alien insects as a driver of biodiversity change are an important yet understudied component of the general threat of biological invasions. The environmental impacts of invasive alien insects are varied and widespread, with evidence to suggest that an insect species global maximum impact is likely to increase in severity as it increases its non-native distribution. Two potential explanations are that large geographic distributions include environmental heterogeneity and increase resource availability, or that there are intrinsic factors of widely-spread species that also facilitate greater impacts. Determining which explanation is more likely, and developing a more comprehensive and general understanding of the environmental impacts of invasive alien species, can be assisted by addressing the information shortfalls highlighted in this research.

## Introduction

Insects are one the largest and most diverse animal groups on land [1]. Unsurprisingly, insects therefore also constitute a large proportion of species with populations established outside of their native range; the total estimated number is as high as one quarter of the potential source pool of alien insect species [2, 3]. Many insect introduction events are unintentional and a consequence of international trade [4]. This is especially true of agricultural and horticultural trade that harbors insects as stowaways, a consequence of the close association between floral commodities and insect occurrence [5, 6]. Not all introduced insect species are, however, expected to negatively affect their recipient environments [7], although the probability that an introduced species will have negative consequences has been shown to increase with the number of introduction events [8, 9]. This propagule pressure is expected to continue for both insects and other groups and understanding the consequences of this process for biodiversity and ecosystems remains key to mitigation and prevention [10].

Alien insects are well-known for their socioeconomic impacts [11], including economic loss [12, 13] from decreased crop yields [14] and management costs [15, 16], through to threatening food security [17] and human health [18, 19]. However, their environmental impacts - realized and potential - are of equal concern. Despite their relatively small size, the cumulative effect of alien insects in a community can cascade through the trophic chain, ultimately affecting environmental processes at an ecosystem scale, mediated via multiple direct and indirect interactions [20, 21]. There are many examples of damaging insect impacts, particularly due to the Hymenoptera (bees, wasps, ants, sawflies [22-26]. A recent review explored the effects of invasive insects on native insect communities [27]. However, generalities on the range and comparative severity of specific environmental impacts (impact mechanisms), and the ways in which these materialize across insect orders, have not yet emerged (although see [28, 29]).

Insects are generally under-represented in biodiversity databases [30] and have taxonomic, distribution and abundance-based knowledge shortfalls [31]. This deficit extends to information on environmental impacts, which are necessarily informed by species population and life history trait information [32]. Beyond a real absence of data (data deficiency), one method for synthesizing information on invasion impacts that does exist, in a useful and targeted form, is environmental impact assessment; a tool used to guide the development of evidence-based risk assessment. Using existing information and a structured protocol to classify the severity of impacts so far realized (i.e., impact history) for alien species improves the potential for predicting future impacts [33], and facilitates triage in species-based prioritization [34]. Evidence-based risk assessments are important for the appropriate prioritization of invasive species for management and prevention [35-37].

Here we take a species-specific approach to assess the environmental impacts of invasive alien insects, at both national and global scales. Our approach was to individually assess and then collate the information across invasive alien insect species, for the purpose of quantifying the geographic and taxonomic variation in the severity and mechanisms by which invasive alien insects impact native environments. We test four hypotheses about invasive alien insect impacts, the first two to better understand and redress bias in evidence and the second two to test biogeographic hypotheses of insect invasion. Specifically, we predict that: (1) There is greater environmental impact evidence for those insects that also have socioeconomic impacts [11, 38]. Species of socioeconomic concern are assumed to receive more research attention, and therefore to have a wealth of information also relevant to their environmental impacts [39]. Support for this hypothesis would demonstrate a bias in knowledge towards species of socioeconomic over environmental concern. (2) The extent of information available to assess the impacts of alien insects differs among insect orders and is biased toward Hymenoptera. This has been suggested previously [25], but quantifying the relative bias across orders here could inform prioritisation of research investment to further quantify environmental impacts. (3) There is a positive relationship between species’ introduced range and environmental impact severity. Geographic range is considered one of three key factors contributing to the impact of an alien species [40]. However, the relationship between invaded range and severity of impact has to date not been tested for any taxon as far as we are aware. A positive relationship between invaded range extent and severity of impact would add further impetus to the priority of preventing invasive alien species spread. (4) The maximum realised impacts of invasive alien insects are greater on islands than mainland locations. For example, the weight of evidence for harmful environmental impacts from alien birds was higher for island-located impacts than impacts than elsewhere [41]. Indeed, for both birds and mammals, islands are well-known to be highly susceptible to negative biodiversity impacts of invasive alien species, however, this has not previously been tested for insects [41, 42].

## Methods

### Species pool assessed

Insect species were extracted from the Global Register of Introduced and Invasive Species (GRIIS) as the initial source pool. GRIIS contains over 2700 introduced and invasive insect species of environmental concern, recorded at a country by species scale [43]. These species were then filtered to the subset identified as damaging to the environment in a range of other databases [28], creating a species pool that can be loosely interpreted as alien insect species of known and potential environmental concern. This resulting pool of 590 insect species, of which 100 high priority species were used in a pilot assessment to test and refine the assessment protocol [28]. These 100 species were re-assessed here so that all species were assessed using the latest information and following exactly the same process. For orders with < 20 species (*n* = 13 orders) in the species pool of 590, all species in the order were assessed. For orders with > 20 species present (*n* = 6), 20 species per order were randomly selected (Table 1). The final set of 352 species assessed across 19 taxonomic orders were thus considered to be taxonomically representative of insect species considered to have negative environmental impacts somewhere in their introduced range.

**Table 1.** Number and percentage of species with environmental impact information was similar overall for species that are and are not considered socioeconomic pests. However, this pattern did not hold for some taxonomic orders. For example, environmental impact information availability differed depending on socioeconomic pest status for Coleoptera and Diptera [110], with non-socioeconomic pests having fewer data deficient (DD) species than pest species. Orders with one species or less across all categories are grouped into the “Other” taxonomic order.

| **Taxonomic Order** | **Data deficient** |  | **Data available** |  |
| --- | --- | --- | --- | --- |
|  | **SE pest** | **Non-SE pest** | **SE pest** | **Non-SE pest** |
| Blattodea | 2 (40%) | 2 (50%) | 3 (60%) | 2 (50%) |
| **Coleoptera** | **24 (75%)** | **8 (47%)** | **8 (25%)** | **9 (53%)** |
| Dermaptera | 7 (88%) | 3 (100%) | 1 (12%) | 0 (0%) |
| **Diptera** | **10 (77%)** | **10 (50%)** | **3 (23%)** | **10 (50%)** |
| Hemiptera | 27 (66%) | 6 (66%) | 14 (34%) | 3 (34%) |
| Hymenoptera | 11 (34%) | 28 (42%) | 21 (66%) | 38 (58%) |
| Lepidoptera | 17 (71%) | 2 (66%) | 7 (29%) | 1 (33%) |
| Psocodea | 2 (100%) | 16 (94%) | 0 (0%) | 1 (6%) |
| Thysanoptera | 8 (89%) | 1 (100%) | 1 (11%) | 0 (0%) |
| Other | 3 (75%) | 24 (86%) | 1 (25%) | 4 (14%) |
| **Total** | **111 (66%)** | **100 (60%)** | **58 (34%)** | **67 (40%)** |

### Impact assessments

Environmental impacts of alien insects were assessed using the Environmental Impact Classification for Alien Taxa (EICAT) protocol [44-46]. EICAT is a semi-quantitative method that uses available evidence to classify the negative environmental impacts of alien species according to their severity and mechanism of impact, i.e., Minimal Concern (MC, impacts on native taxa negligible), Minor (MN, no evidence for a decline in population sizes of native taxa), Moderate (MO, impact native species population sizes but no evidence of local apparent extinction), Major (MR, reversible local extinction of one or more native taxa) and Massive (MV, irreversible local extinction of one or more native taxa) (for full descriptions of categories used see [28]). Typically, the reported impact severity (and associated mechanism and confidence) for a given alien species is the global maximum impact severity recorded. However, this can omit useful and informative information pertaining to variation in both how and where alien species are negatively affecting the environment. As such, in this study a focus on the global maximum recorded impact severity only occurred when examining the relationship between alien geographic range and impact severity. All other descriptive and statistical analyses are based on all recorded impact evidence where relevant, i.e. including multiple, independent pieces of evidence for multiple species.

Web of Science (WoS, www.webofknowledge.com) was the primary database used for literature searches. Differential literature use, for example due to different search strategies, is a primary reason for assessment outcomes differing among independent assessors [28]. Therefore, taking a systematic approach to searching available impact evidence decreases uncertainty in the assessment outcome. Literature searches occurred using *All Databases*, as opposed to *Web of Science Core Collection*, with the timespan selected as *All years*. For a given species, the search string consisted of the species binomial (or trinomial in the case of subspecies) name with the Boolean operator OR used to separate all known synonyms. Accepted names and synonyms were determined using the Global Biodiversity Information Facility (GBIF) taxonomic backbone. Using only species names can result in a large number of search returns. Although this leads to the return of many irrelevant publications, consequently increasing the time taken to complete an assessment, it also decreases the likelihood of missing relevant information that contains a more nuanced form of impact evidence. Additionally, this species-specific approach to searching the literature enables a more complete picture of the state of alien insects and their environmental impacts, as much impact research is biased toward a subset of well-studied species [25, 32, 47]. Publications warranted examination if the title and/or abstract provided any indication that the alien insect species in question was negatively affecting the native environment in a location outside of the species native range. Publications that were deemed potentially relevant based on their title that were not published in English were translated using Google Translate to confirm degree of relevance and avoid English-biased research [48]. Following the WoS searches, the same search strings were included in Google Scholar with the first 50 returns checked for titles not returned in the WoS search. To ensure transparency, all literature search information, including dates of search, terms used, and the search outcomes, were recorded and is available online, along with all code for replicating the analyses (https://doi.org/10.5281/zenodo.6637035).

Complementing the literature searches were two other secondary sources of information. First, potentially relevant studies that were cited by an examined paper but were not themselves included in the literature search returns were also assessed. Second, Centre for Agriculture and Bioscience International (CABI) species datasheets from both the Crop Pest Compendium (CPC) and Invasive Species Compendium (ISC) were examined for any additional information on each species. The presence or absence of both CPC (pest) and ISC (invasive species) datasheets were also recorded for testing the hypothesis that socio-economic pests are more likely to have evidence of environmental impact.

Studies included for assessment were those in which there was evidence of an *introduced insect species negatively affecting the native biodiversity/environment*. An example of a study type *not* included is one in which native species were negatively affected by human actions related to management of the alien species under assessment, e.g., a chemical spray intended for the alien species that negatively affects native species. Similarly, if humans introduced a biocontrol agent to control the alien species under assessment, and that biocontrol agent negatively affects the native environment, this was also not considered (it would, however, be evidence under an assessment of the biocontrol agent itself). Furthermore, the focus was on *what* was affected, not *where* it was affected. For example, whilst evidence of an alien insect damaging an agricultural crop was not included, evidence for the same insect negatively affecting a native species *within* an agricultural crop was included.

### Data analysis

Species were classed as a known socioeconomic pest if they had “pest” listed on their CABI CPC datasheet; datasheets were used as a proxy for socioeconomic pest status. If a species did not have “pest” listed on their CABI CPC datasheet, or a datasheet was absent, they were not considered a socioeconomic pest. As such, this was treated as a binary outcome variable (yes/no) and was assessed against the information availability binary variable (Data Deficient/non-Data Deficient). Log-linear analysis was used to test the hypothesis that known socioeconomic pests are more likely to have evidence of environmental impact. Additionally, a Wilcoxon rank sum test was performed to compare the number of WoS search returns between species that were, and were not, considered to be socioeconomic pests.

Log-linear analysis was used to test the hypothesis that the extent of environmental impact information, i.e., the proportion of non-data deficient species, differed among taxonomic orders. Information availability was considered a binary outcome variable (data deficient/non-data deficient) and the expected number of each outcome instance across all 19 taxonomic orders assessed was analysed.

Assessing the relationship between impact severity and alien geographic range would ideally use extent of occurrence (EOO) [49] as the measure of range size. However, given the taxonomic and spatial biases present in species occurrence data [30, 50], we chose instead to use the number of countries listed for a species in the Global Register of Introduced and Invasive Species (GRIIS), which includes verified records of presence at a country scale, as a proxy for alien geographic range [43, 51]. Ordinal logistic regression was used to test the hypothesis that impact severity increases with non-native geographic range size. Only species with evidence of impact were used in the analysis (i.e., excluding any Data Deficient species), with global maximum impact severity and the number of countries with alien populations as the response and explanatory variable, respectively.

Under the EICAT framework, impact severities of Moderate or higher are considered “harmful” [46]. To test the hypothesis that islands experience a higher proportion of harmful impacts from invasive alien species than mainland locations, log-linear analysis was used, treating impact severity as a binary outcome variable (harmful/not harmful) and assessing it against the landmass type binary variable (island/mainland).

## Results

### Pest status and data availability

When all species were pooled there was a significant difference in WoS search returns based on socioeconomic pest status, with those considered socioeconomic pests returning a higher median number of search returns (*U* = 9069, *n*_*1*_ = 180, *n*_*2*_ = 172, *p* < 0.001, Figure S1). The effect size for this difference was *r* = 0.33, indicating an intermediate effect (Cohen 2013) of socioeconomic pest status on WoS search returns. Although there was a significant difference between the number of species Data Deficient and non-Data Deficient for environmental impact information, this difference was independent of socioeconomic pest status (χ^2^ = 0.012, *df* = 1, *P* = 0.9131). The number and proportion of Data Deficient species was similar between the two pest status groups (Table 1). For specific insect orders, the general pattern of a higher proportion of Data Deficient species remained, but with some exceptions (Table 1). Although Coleopteran and Dipteran that are socioeconomic pests were more commonly Data Deficient for environmental impact information, for species not considered socioeconomic pests the proportions were either equal (Diptera) or higher for species with evidence of impact (Coleoptera) (Table 1). Therefore, although overall socioeconomic pest status shows no association with environmental impact information availability, for two comparatively species rich orders the pattern was counter to expected, with less evidence available on the environment impacts of socioeconomic pests than for species that do not have socioeconomic impacts.

There was strong evidence of an association between taxonomic order and the number of species assessed as Data Deficient. The probability that the independence model performed as well as the model containing an interaction between taxonomic order and assessment outcome (DD vs Not DD), if there was no association, was small (χ^2^ = 69. 834, *df* = 17, *P* < 0.0001). For most orders, the proportion of Data Deficient species is greater than non-Data Deficient for evidence of environmental impact. Exceptions to this were the Blattodea, Hymenoptera, and Mantodea (Figure S2, Table S1). There was, however, high variation in the proportion of species assessed as Data Deficient across orders – ranging from 0 to 1 (Figure S2).

Evidence of environmental impact was available for 125 species, with approximately two thirds (*n* = 211; 59.9%) of species assessed as Data Deficient (Table 2). Quantity of impact evidence varied widely, with a mean of 4.9 ± 7.1 SD impact publications per species (Figure S3, Figure S4). The relatively high standard deviation reflects the presence of a small number of disproportionately well studied species (*Harmonia axyridis, Philornis downsi, Adelges tsugae, Apis mellifera, Bombus terrestris, Linepithema humile, and Solenopsis invicta*) with 15 to 50 publications demonstrating evidence of impact (Figure S3, Table S2). Similarly, the number of species with evidence of impact was disproportionately higher in North America and Australia than elsewhere (Figure 1A). Of the species assessed, Hymenoptera had most impact evidence (*n* = 59) followed by Coleoptera (*n* = 17), (Table 2). There was no evidence of populations established outside their native range for 16 of the 352 species assessed (subsequently denoted NA).

**Table 2.** The number of species assessed in each insect order and the percentage for which at least one item of evidence on environmental impact was found. Parentheses include percentages of species assessed from species pool (Assessed), and species assessed with impact evidence (Evidence). Although three species of Trichoptera were initially included from the species pool, all three were concluded to have no populations established outside their native range (i.e. NA).

| **Order** | **Assessed** | **Evidence** | **Species pool** |
| --- | --- | --- | --- |
| Hymenoptera | 98 (77%) | 59 (60%) | 128 |
| Coleoptera | 50 (34%) | 17 (34%) | 146 |
| Hemiptera | 49 (60%) | 16 (33%) | 81 |
| Diptera | 35 (80%) | 14 (40%) | 44 |
| Lepidoptera | 31 (55%) | 8 (26%) | 56 |
| Blattodea | 9 (100%) | 5 (56%) | 9 |
| Mantodea | 2 (100%) | 2 (100%) | 2 |
| Siphonaptera | 4 (100%) | 1 (25%) | 4 |
| Thysanoptera | 10 (100%) | 1 (10%) | 10 |
| Dermaptera | 17 (100%) | 1 (6%) | 17 |
| Psocodea | 20 (33%) | 1 (5%) | 66 |
| Orthoptera | 2 (100%) | 0 (0%) | 2 |
| Phasmatodea | 6 (100%) | 0 (0%) | 6 |
| Zygentoma | 6 (100%) | 0 (0%) | 6 |
| Ephemeroptera | 2 (100%) | 0 (0%) | 2 |
| Odonata | 1 (100%) | 0 (0%) | 1 |
| Embioptera | 5 (100%) | 0 (0%) | 5 |
| Neuroptera | 2 (100%) | 0 (0%) | 2 |
| Trichoptera | 3 (NA) | 0 (NA) | 3 |
| **Total** | **352** | **125** | **590** |

**Figure 1.**
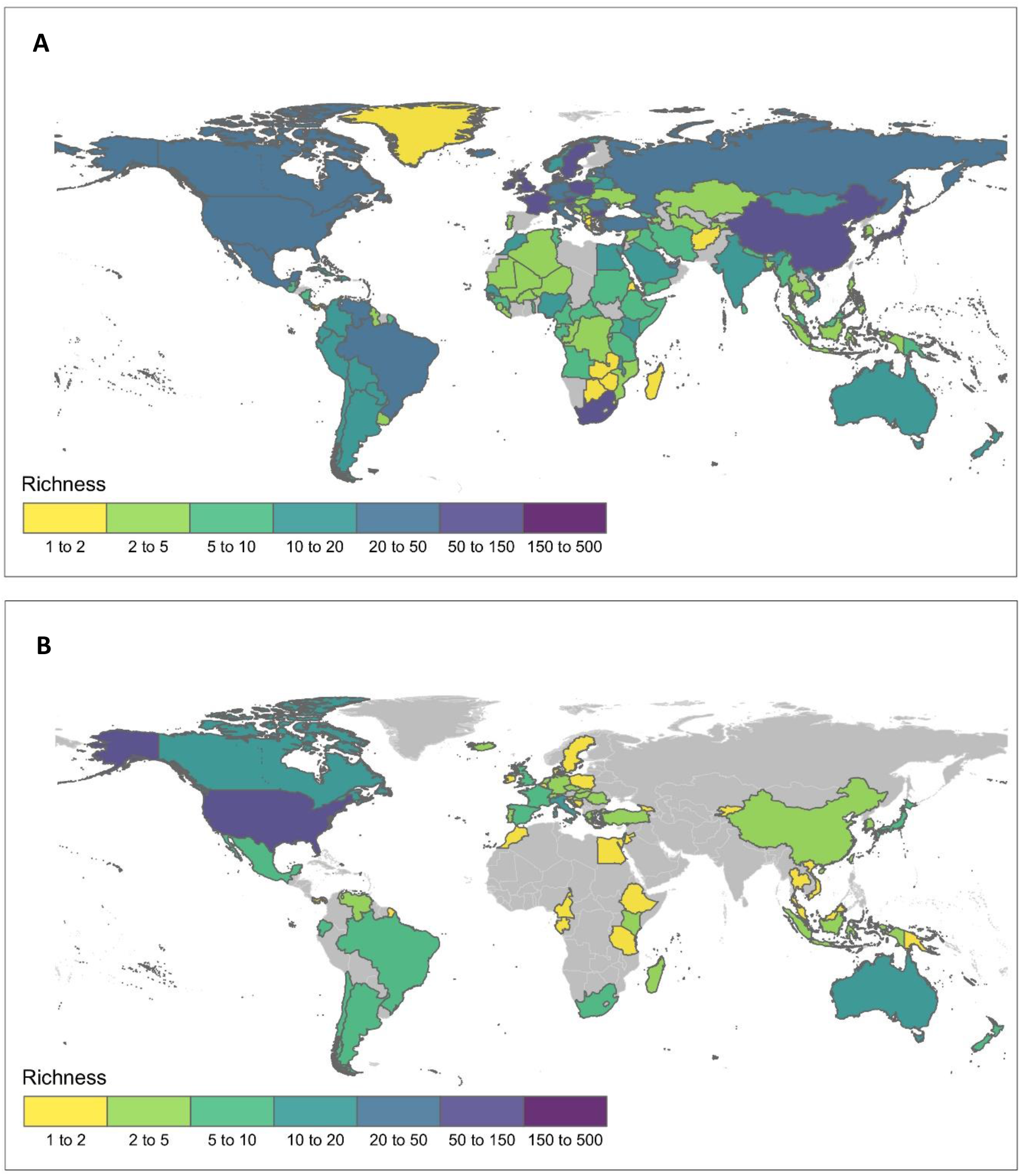
A. Global distribution of known alien and invasive insect populations that harm the environment at a national scale (*n* = 7049, according to GRIIS, as of 11^th^ June 2020). B. Number of alien insect species with published evidence of environmental impact per country. Country boundary data was obtained from the Global Administrative Areas (GADM) database [111].

### Impact mechanism, severity and confidence

Evidence of environmental impact was found on 11 impact mechanisms. Competition was the most frequently attributed impact mechanism across all instances of impact evidence (*n* = 244), followed by herbivory (*n* = 135), predation (*n* = 95), and parasitism (*n* = 35) (note a species may have evidence of impact for multiple mechanisms). As expected, dominant mechanisms differed across orders; for example, all cases of environmental impact via hybridization were associated with Hymenoptera, although overall hybridization was not a common mechanism amongst mechanisms for Hymenoptera (Figure 2, Figure S5, Figure S6). Similarly, although transmission of disease was most frequently attributed to Coleoptera, disease transmission was associated with only four species (Figure 2, Figure S5, Figure S6).

**Figure 2.**
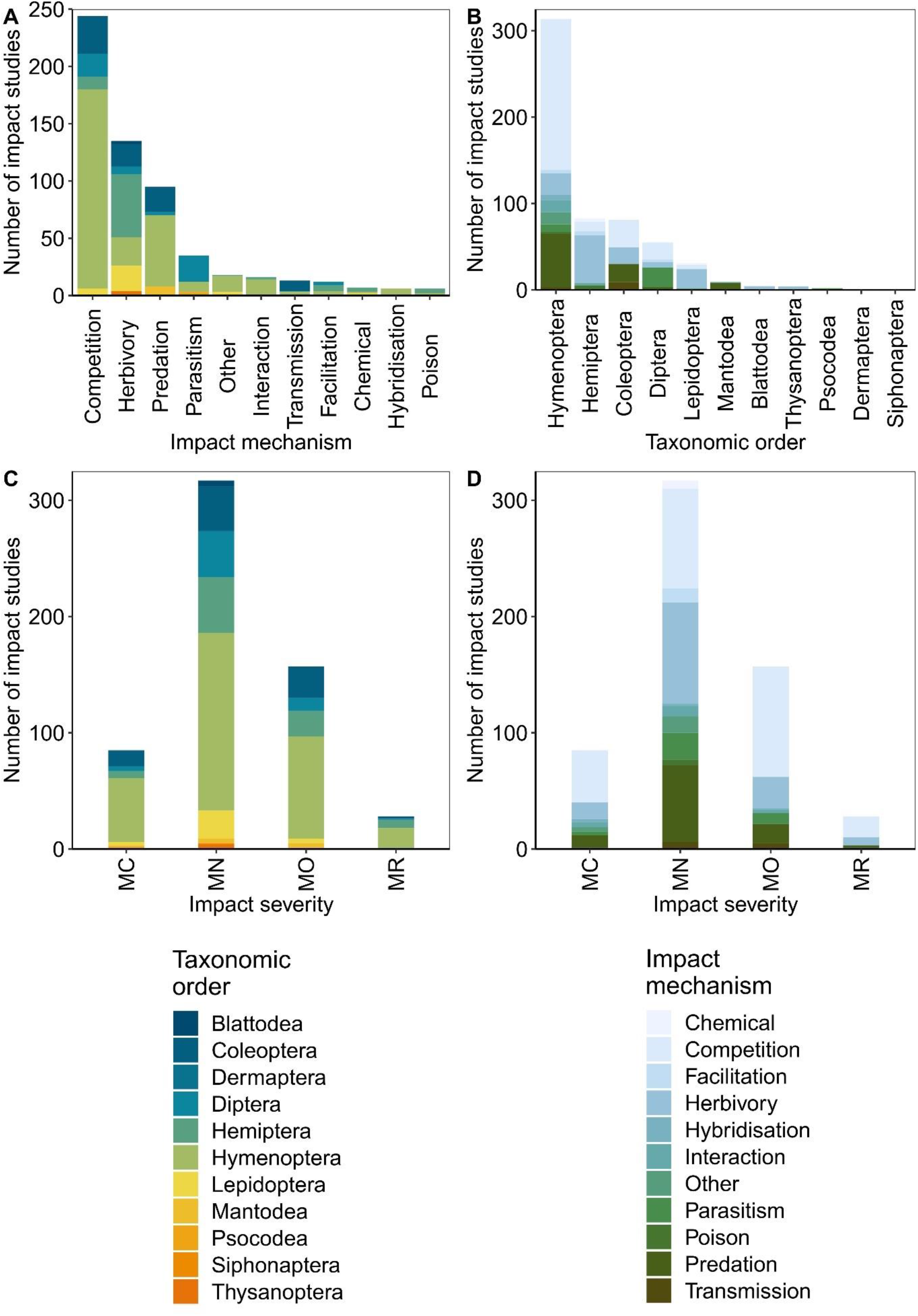
Occurrence frequencies for different combinations of impact mechanism, severity and taxonomic order. Each plot represents the number of impact studies for: A. each environmental impact mechanism, per taxonomic order, B. each taxonomic order, per impact mechanism, C. each impact severity category, per taxonomic order and D. each impact severity category, per impact mechanism. Impact severity categories are Minimal Concern (MC, impacts on native taxa negligible), Minor (MN, no evidence for a decline in population sizes of native taxa), Moderate (MO, impact native species population sizes but no evidence of local apparent extinction), Major (MR, reversible local extinction of one or more native taxa) (there were no species assessed as having a Massive (MV) impact, irreversible local extinction of one or more native taxa) (for full descriptions of categories used see Clarke et al. (2021)).

The most common impact severity category attributed to species across all impact evidence was Minor (*n* = 317). This was followed by Moderate (*n* = 157), Minimal Concern (*n* = 85), and Major (*n* = 28), with no instance of a Massive impact (irreversible extinction) being attributed. Hymenoptera had the highest proportion of recorded evidence for each category of impact severity (Figure 2, Figure S5, Figure S6). However, variability in impact severity was highest in the Hymenoptera (Figure 2, Figure S5, Figure S6). The largest proportion of impact severity classifications for all taxonomic orders was Minor (except Mantodea), and no species of Blattodea, Dermaptera, Siphonaptera, and Thysanoptera had impacts in higher severity categories.

Confidence ratings for each assessment of an individual piece of impact evidence varied, with Low confidence the most commonly attributed rating (*n* = 349), which was followed by Medium (*n* = 284) and High (*n* = 234). Little to no correlation existed among the variables of taxonomy, impact mechanism, impact severity and confidence, with most points located around the origin of the first two dimensions which collectively only explained 14.4% of the variation (Figure 3).

**Figure 3.**
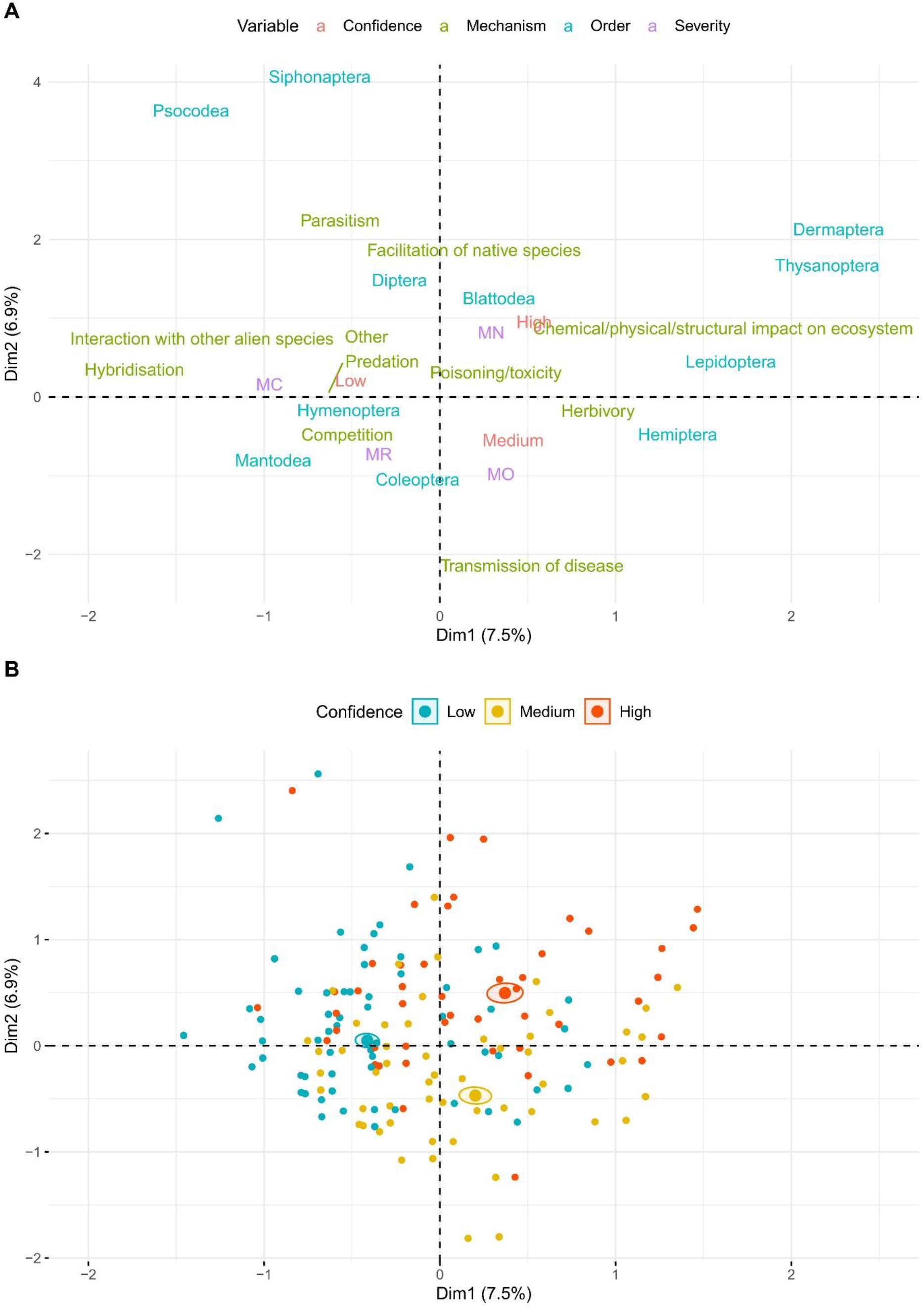
A. Biplot showing the first two dimensions of a multiple correspondence analysis (MCA) examining potential correlations among insect order, impact mechanism, impact severity, and confidence rating (Low, Medium, High) for each instance of environmental impact evidence. Levels of each categorical variable are plotted, with distance between each variable level corresponding to similarity in profiles. Little to no correlation exists among the four variables, with the exception of some expected grouping such as herbivory and Hemiptera or competition and Hymenoptera. Results further support the claim for the context dependence of environmental impacts. B. The distribution of confidence ratings of each impact study, with colour reflecting Low, Medium, or High confidence. The ellipses represent uncertainty of the mean coordinate point for each confidence category. For example, the Low confidence ellipse is smaller than the other ellipses which can be seen as the smaller spread of Low confidence studies (blue dots) across the first two dimensions. Impact severities are: Minimal Concern (MC), Minor (MN), Moderate (MO) and Major (MR).

### Geographic range and impact severity

Alien insect populations of species qualitatively considered to be of some or potential environmental concern (*n* = 590) are known from most countries of the world (Figure 1A). In total there were 864 species-country combinations. Countries in Asia, Western Europe and Southern Africa have the most established alien insect species in this pool of species that represents the majority of alien insects of environmental concern. The geographic distribution of populations with evidence of impact is patchier (Figure 1B). Alien geographic range (using number of country records per species in GRIIS) was significantly associated with maximum impact severity (χ^2^ = 8.43, *df* = 1, *P* = 0.004). Predictions show that for a given insect species, the probability that its maximum impact severity will be Major increases as the number of countries with established populations increases (Figure 4). For every one-unit increase in alien geographic range, the odds of the maximum impact being more severe are multiplied by 1.02, i.e., a two percent increase (95% CI 1.01, 1.03). In contrast, the probability that its maximum impact severity will be Minimal Concern or Minor decreases with the number of countries in which it becomes established (Figure 4). The probability that the maximum impact severity of a species will be Moderate remains approximately constant, regardless of the number of countries in which it is established, a result of the high variation in geographic range (Figure 4).

**Figure 4.**
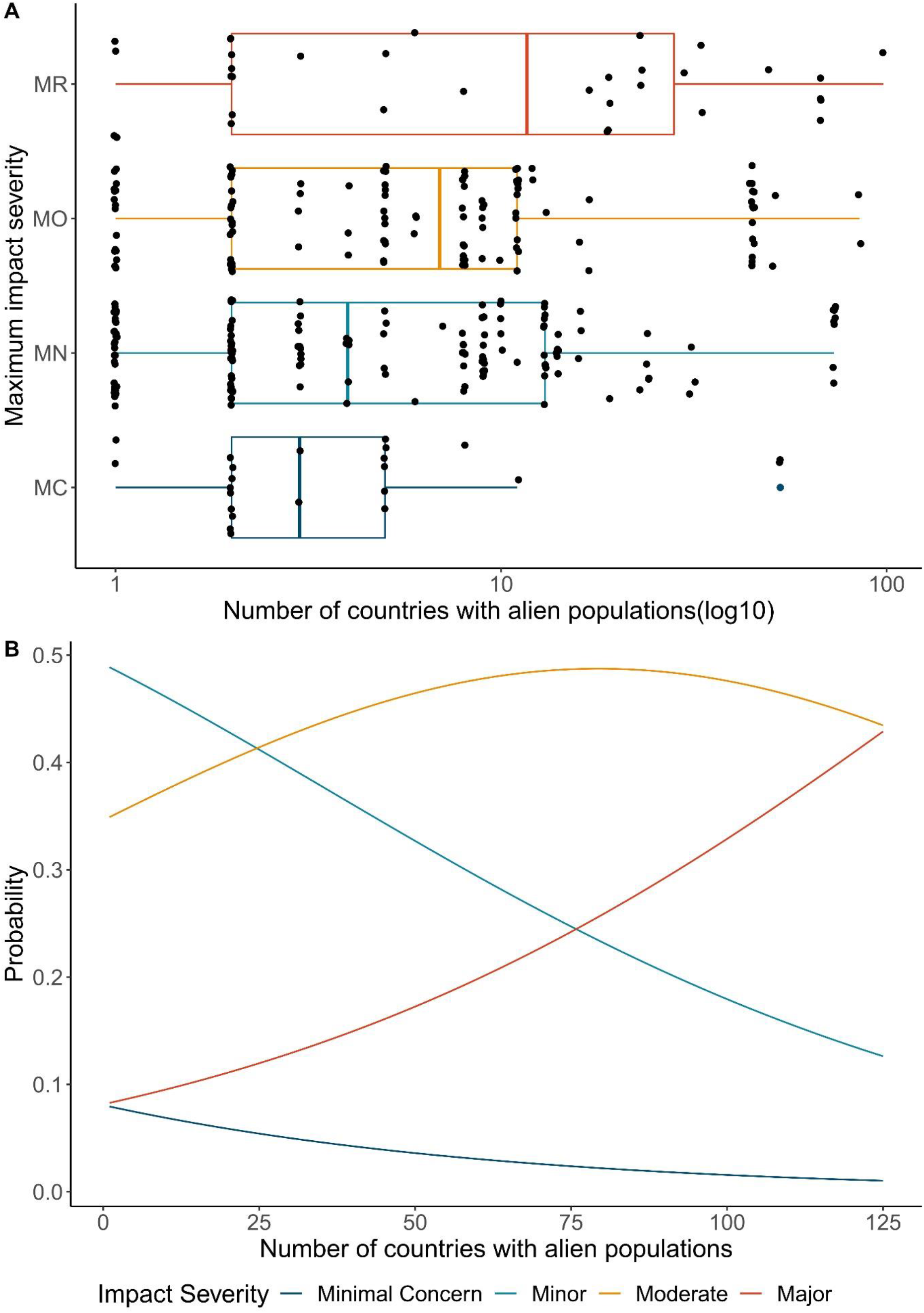
Association between geographic range and environmental impact severity of alien insects. A. Global maximum impact severity for invasive alien insects is a function of alien geographic range. The probability of an alien insect species global maximum impact being more severe increases with the number of countries in which it has established. Likewise, the probability of less severe maximum impacts occurring decreases with geographic range. B. Alien geographic range variation (number of countries with alien populations) for species within each maximum impact severity classification. Most species had a maximum impact of Minor or Moderate severity. Impact severity categories are Minimal Concern (MC, impacts on native taxa negligible), Minor (MN, decreased performance of native taxa but no evidence of population decline), Moderate (MO, native species population declines but no evidence of local apparent extinction), Major (MR, reversible local extinction of native taxa) (there were no species assessed as having a Massive (MV) impact, irreversible local extinction of one or more native taxa) (for full descriptions of categories used see [28].

### Islands and impact severity

No evidence of an association between landmass type (continent vs island) and impact severity (harmful vs not harmful) was found, with a well-fitting independence model (χ^2^ = 0.154, *df* = 1, *P* = 0.901). Although the proportion of harmful to not harmful species was equivalent across landmass types (Table 3), landmass as a variable was retained, as including impact severity as a sole predictor variable led to an inadequate model (χ^2^ = 101.581, *df* = 2, *P* < 0.0001). Importantly, this analysis was based on each unique impact study and does not account for any differences among taxonomic order or insect species.

**Table 3.** Summary information comparing harmful environmental impact information between island and mainland landmasses. Harmful impacts are those characterized as impact severities that range from being responsible for population declines to irreversible extinction of native species (i.e. include Moderate (MO), Major (MR) and Massive (MV)). There was no difference in the proportion of impact studies that had evidence of harmful impacts between island and mainland landmasses. Information was based on all environmental impact evidence, i.e., some species were included more than once.

|  | **Island** | **Mainland** |
| --- | --- | --- |
| Total number of species | 53 | 94 |
| Total number of impact studies | 154 | 384 |
| Impact severity frequency:<br>MC, MN, MO, MR, MV | 23, 78, 46, 7, 0 | 54, 200, 110, 20, 0 |
| No. harmful impacts (percentage of total) | 53 (34%) | 130 (34%) |
| Number of studies per species (mean) | 3.28 ($\sigma^2 = 4.00$) | 4.59 ($\sigma^2 = 7.40$) |
| Number of studies per species (median) | 1 | 3 |

## Discussion

Invasion ecology is replete with hypotheses for explaining various aspects of biological invasions, but few relate to impact events at a global scale [52]. Here we assessed the environmental impact severities and mechanisms of impact for the 125 best-known invasive alien insect’s species globally to assess the availability of impact evidence for this group and to test hypotheses about the geographic distribution of their impacts. Evaluation of a further ∼250 species revealed current evidence is not available to assign environmental impacts. In addition to a highly uneven spread of knowledge across these taxa, we found mixed support for hypotheses associated with invasive alien species [53] and their impacts suggesting, inter alia, that generalities in the environmental impacts of invasive alien insects may differ in some respects from other, better known taxonomic groups of IAS.

### Geographic determinants of environmental impact

Knowledge of the extent to which alien insects have an impact throughout their geographic range is important to acquire for prioritizing invasive alien species, because alien insect species do not necessarily have the same severity of impact throughout their introduced range (e.g., *Pheidole megacephala, Anoplolepsis gracilipes*). However, we found that as the number of countries an alien insect species was introduced to increased, so too did the probability that it would have a maximum global impact leading to local native species extinction. Two potential processes may explain this finding: 1) the broader the global introduced geographic range of a species, the more likely it will have a severe impact somewhere within that introduced range (e.g., empty niche [54], opportunity windows [55]), or 2) species with severe impacts have (on average) larger invaded ranges because of some intrinsic biological property (e.g., ideal weed [56, 57]). More species-specific impact research, within and across taxonomic groups, is required to determine which, if either, of these hypotheses holds.

The commutative property of the three variables that together determine the size of the impact of an invasive alien species – geographic range (area of occupancy), abundance, and per capita or per area effect - mean that the environmental impact of a species could be largely a function of a species abundance and/or per capita effect [40, 58]. For example, the parasitic larvae of the fly *Philornis downsi* are largely responsible for the population decline of multiple endemic bird species in the Galapagos archipelago [59]. It is potentially causing irreparable damage, yet it is only known to have alien populations in two locations, the Galapagos and Brazil [59]. In fact, the probability that an insect species maximum impact would involve the decline of native species populations (i.e., classified as Moderate impact) remained approximately constant with an increasing geographic range. Therefore, although the likelihood of maximum impact severity increases with geographic range, a widespread alien insect’s range does not necessarily predicate severe environmental impacts.

We found no evidence for the island susceptibility hypothesis for invasive alien insects. Island ecosystems were no more likely to suffer from the harmful impacts of invasive insects than mainland locations. In fact, the proportion of impact studies that recorded evidence of a harmful impact was the same for the two landmass types. While our results are congruent with previous research on the environmental impacts of invasive alien insects that likewise determined islands to be no more likely to experience severe impacts than mainland locations [47], our work represents a more taxonomically and geographically comprehensive assessment of the island susceptibility hypothesis. Islands have generally been identified as hotspots for IAS mediated extinctions [42, 60-63], and represent one of the more studied ecosystem types for the environmental impacts of IAS [32]. Indeed, at least seven insect species assessed here have caused severe impacts on islands around the globe, including the local extinction of native species [64] and changes to entire ecosystem structure [65].

### Mechanisms of environmental impact

Competitive interactions were the most prevalent form of environmental impact. This was particularly prominent within the Hymenoptera, which had a disproportionately higher amount of competition-driven negative impacts than other taxonomic orders. Hymenoptera were found to outcompete natives via both interference [66, 67] and exploitative [68, 69] competition, and for some species (*Anoplolepsis gracilipes, Linepithema humile, Myrmica rubra, Pachycondyla chinensis, Paratrechina longicornis, Pheidole megacephala, Solenopsis invicta, Apis mellifera, Apis mellifera scutellata*), outcompeting natives was the mechanism responsible for their most severe impacts in which some native species were driven to local extinction [53, 65, 70-76]. Only two non-Hymenopteran species (*Chrysomya albiceps* (Calliphoridae, Diptera) and *Digitonthophagus gazella* (Scarabaeidae, Coleoptera)) caused as severe impacts through competition [77, 78]. For orders other than Coleoptera, competition was not the most prevalent mechanism of impact.

Competition as the dominant driver of severe environmental impacts in insects differs from the finding that predation is the mechanism by which other animal taxa, especially mammals and fish) bring about declines in native species [42, 79]. Nonetheless, consumer-resource interactions, such as predation and herbivory were the next most prevalent and most severe impact mechanisms for invasive insects after competition. For example, by preying upon a native wingless fly, the carabid beetle *Merizodus soledadinus* is likely responsible for the fly’s local extinction on the Kerguelen Islands [80]. Similarly, the big-headed ant (*Pheidole megacephala*) and the little fire ant (*Wasmannia auropunctata*) are both recorded as preying upon native species to the point of local extinction in Hawaii [64] and Gabon [81], respectively. Herbivorous insects have likewise caused severe environmental impacts, the most notable of which is the hemlock woolly adelgid (*Adelges tsugae*) whose damage has led to large sections of eastern hemlock mortality on the eastern coast of the United States [82]. Two other species, the cottony cushion scale (*Icerya purchasi*) and the cycad aulacaspis scale (*Aulacaspis yasumatsui*), are similarly responsible for severe herbivorous impacts on native flora in Guam and the Galapagos Archipelago [83, 84].

The Hymenoptera represented the taxonomic order with the largest proportion of recorded impacts, in terms of total number of species with evidence and total number of instances of impact evidence. However, the results suggest that this is may be a function of the bias toward studying this group of insects, which contains some of the most well studied invasive insects such as the red imported fire ant and the Argentine ant [25, 32]. To date, species of Blattodea, Dermaptera, Siphonaptera, and Thysanoptera have been recorded as having only minor or minimal environmental impact. There are two reasons why a species’ maximum impact severity is low; either (i) there is an absence of evidence for more severe impacts, or (ii) there is evidence that the species is not having more severe impacts, i.e. evidence of an absence of an effect [28]. Most often it is the former rather than the latter, suggesting that a precautionary approach would be wise until evidence of an absence of impact has been gathered for each species.

### Why are there so many data deficient species?

Geographical biases in IAS impact research are well known [25, 32, 85, 86]. Here, for example, of the 864 species-country insect impact combinations, 59% occurred within the Americas IPBES region, of which 83% occurred within North America. Indeed, the IPBES subregions of North America, Western Europe, and Oceania accounted for 75% of all recorded environmental impact evidence. However, countries within these subregions also generally have higher numbers of recorded insect introductions. Nevertheless, even countries within these subregions with a relatively large number of known alien insects have a pronounced impact research deficit [85]. For example, Denmark contains at least 43 insect species recorded as introduced, but only one (*Harmonia axyridis*) was found to have evidence of environmental impact in that country [87, 88]. Similarly, many species only have impact evidence from a small subset of countries within their introduced range. For example, *H. axyridis* is one of the most widespread insect species, with introduced populations in at least 45 countries, has evidence of environmental impact in only 14 (∼ 30%). That is, there is a large research deficit from both a country and species perspective when examining the environmental impacts of alien insects [85].

Given the many known alien insect populations worldwide, and the importance of understanding and managing the environmental impacts caused by these populations, the question arises as to why in particular there is such a dearth of environmental impact information available? Evans et al. [89] concluded that environmental impact knowledge gaps for alien bird populations were not randomly distributed. Factors such as short residence time, small relative brain size and small alien geographic range were identified as being associated with data deficient species [89]. Here, similar non-random distributions were identified for insects, with the proportion of species classed as Data Deficient differing according to taxonomic order. All species within the orders Embioptera, Ephemeroptera, Neuroptera, Odonata, Orthoptera, Phasmatodea, and Zygentoma were classed as Data Deficient, and only three orders, Blattodea, Hymenoptera, and Mantodea had more species with evidence of impact than without.

Three general reasons are proposed for why a given alien species may be lacking information on environmental impacts. The first is a general lack of interest in a species or that it is perceived as not enough of an environmental threat to warrant concern and subsequent research [86, 89]. Research interest in a species is somewhat dictated by the public’s awareness and interest in that species [90]. As such, species not placed on official IAS lists are likely to be overlooked in a management context [91]. Another possible reason is a species status as a known socioeconomic pest [11]. Impacts such as damage to food crops leading to decreased yields and monetary losses [12, 14, 17] and the spread of disease [19] capture the attention of the public and industry and tend, therefore, to be better funded than environmental impact research. For example, the socioeconomic impacts of invasive alien gastropods are preferentially studied over their environmental impacts [92]. However, although such socioeconomic pests attract more research in general [39, 93], we found no difference in the availability of environmental impact evidence for those species with, versus without, socioeconomic impact. Finally, the perception of a species as an environmental threat results in management interventions that prevent species impacts from being realized. Prevention is the preferred IAS management approach as the feasibility for successful eradication decreases with an increase in the range or population size of an introduced species [94-96]. For example, the citrus long-horned beetle (*Anoplophora chinensis*), primarily known for its effects on fruit trees [97], is also perceived as an environmental threat to native species yet was classed as data deficient due to the perceived threat warranting enough concern for intervention. As such, for some species there may be a discrepancy between the realized and potential impact severity, with a high potential sometimes leading to a smaller realized impact than expected.

It is important to emphasize that impact assessment outcomes require careful interpretation, noting that they are specific to certain situations. For example, if a given insect species is assessed as having a Moderate impact, this applies to a specific alien population, within a specific spatial and temporal context and often is only referring to a single native species being affected. One example of this is the gypsy moth (*Lymantria dispar*) where, depending on impact location and the species affected, has had a negative, neutral, or positive effect on native species [98, 99]. Additionally, meaningful comparisons of assessment results among insect species is made difficult by the high variability in study designs used to record environmental impact evidence [100]. One aspect of this relates to study duration which can vary greatly (e.g., months to years). The importance, and dearth, of studies examining the long-term impacts of invasive species has also been identified more generally [101]. Related to long-term observations, boom-bust dynamics are one example of a phenomenon that can influence the ultimate impact of an invasive species on its recipient environment, as well as influencing potential management strategies [102]. Such dynamics have been observed in insects, particularly in invasive ants [103-106]. Another consideration is the different approaches researchers have employed to determine the effect of invasive species on the native environment. Appropriate comparisons of environmental impacts require the use of a common study design [100]. One common method is to compare invaded and uninvaded sites, inferring impact from negative species co-occurrences. Depending on the design, this can potentially overestimate impact severity by treating species absences as evidence of mortality as opposed to emigration [107]. Regardless, it has been argued that such information alone cannot conclusively lead one to infer invasive alien species as the proximal cause of the observed negative co-occurrence [108], where the IAS could simply be a passenger of change rather than a cause [109].

## Conclusions

Here we show that the impacts of invasive alien insects are, on average, not more severe on islands than on the mainland, and that competition, followed by cross-trophic level interactions, (as in other animal groups), are more frequently associated with the most severe environmental impacts in alien insects. The hypothesis, and the processes underpinning it, that the probability of a population having a Major environmental impact at any particular locality increases with the number of localities in which it has established outside its native range (in parallel with the propagule pressure hypothesis, but further along the invasion continuum) requires further examination, both within and across taxonomic groups. Regardless, this finding provides further impetus for the importance of prevention and limiting the spread of invasive alien species. Finally, we have further demonstrated the geographic and taxonomic biases present in our knowledge of environmental impacts of invasive species. The ability to develop predictions and generalize results requires an even spread of knowledge across species, space and time. A more strategic approach to generating evidence of environmental impact would benefit national and local management scale prioritization for invasive alien insects, targeting priority information gaps such as underexplored species and areas, and testing theory to identify generalities.

## Conflict of interest statement

The authors declare no conflicts of interest.

## Acknowledgements

DAC acknowledges support from an Australian Government Research Training Program (RTP) scholarship. DAC and MAM acknowledge support from the ARC SRIEAS Grant SR200100005 Securing Antarctica’s Environmental Future.

## Notes

https://doi.org/10.5281/zenodo.6637035

### Competing Interest Statement

The authors have declared no competing interest.

