## Supplementary material for "Invasive alien insects represent a clear but variable threat to biodiversity"

Supplementary information

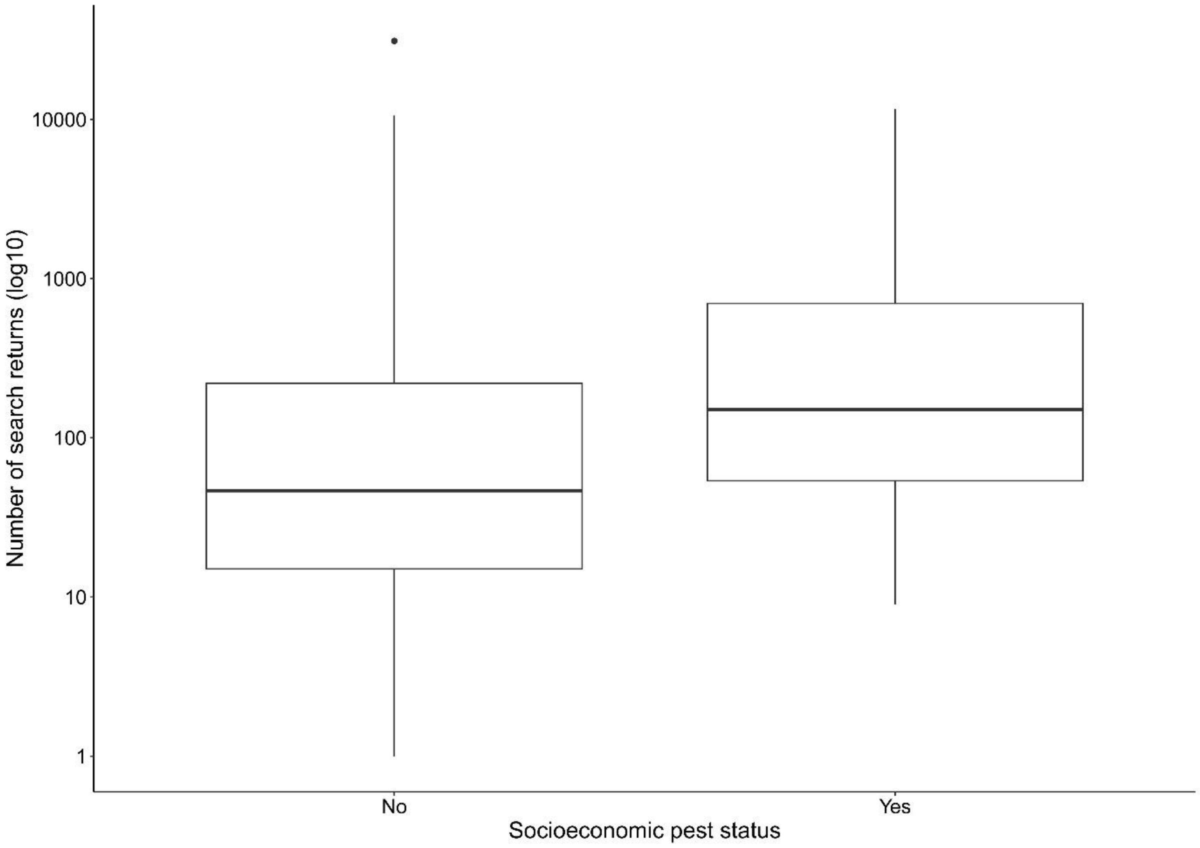


Figure S1. Number of Web of Science literature search returns (log10) for alien insect species, grouped according to their status as known socioeconomic pests. Insect species considered to be of socioeconomic concern had a significantly higher number of literature search returns, indicating a potential research bias toward this group of species.

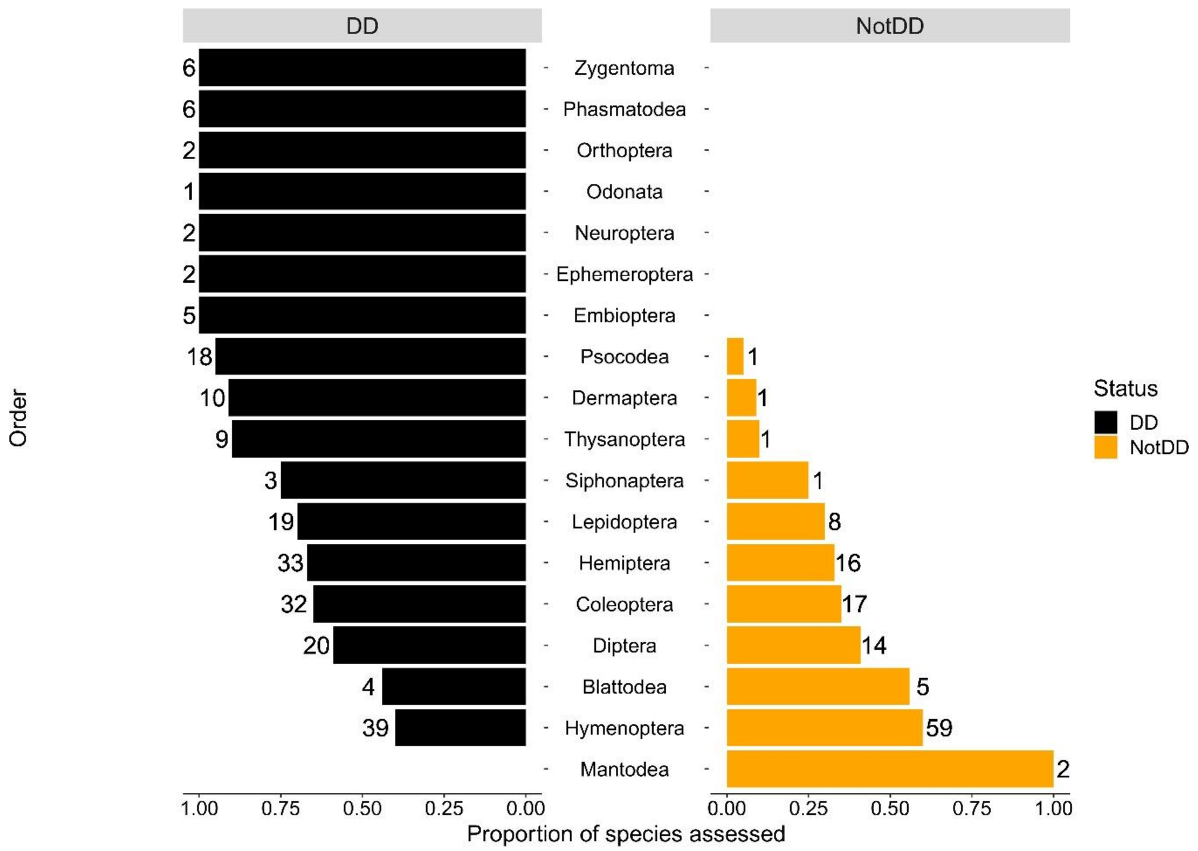


Figure S2. For each taxonomic order, the proportion (and number) of species assessed that were determined to be data deficient (DD) and non-data deficient (NotDD) for environmental impact information.

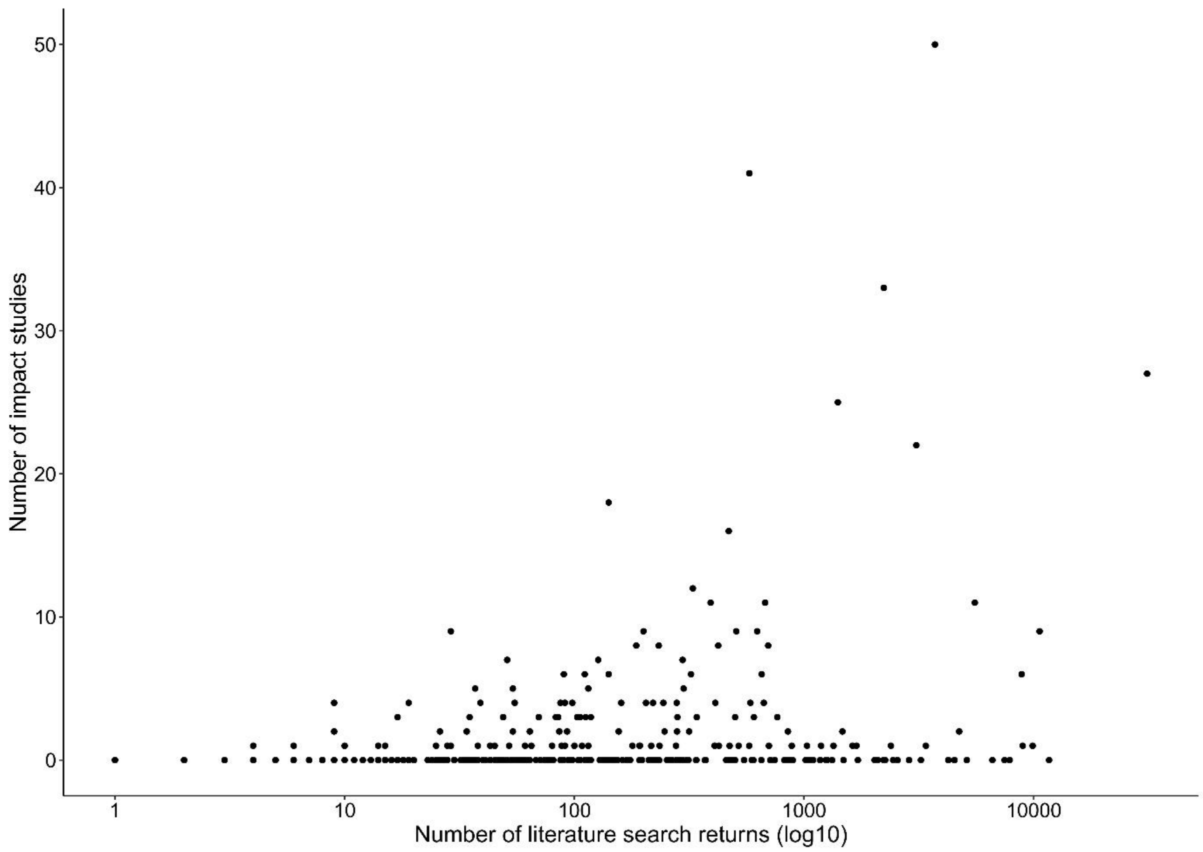


Figure S3. Relationship between the number of Web of Science literature search returns (log10) for each alien insect species assessed and the number of studies examining their negative environmental impacts.

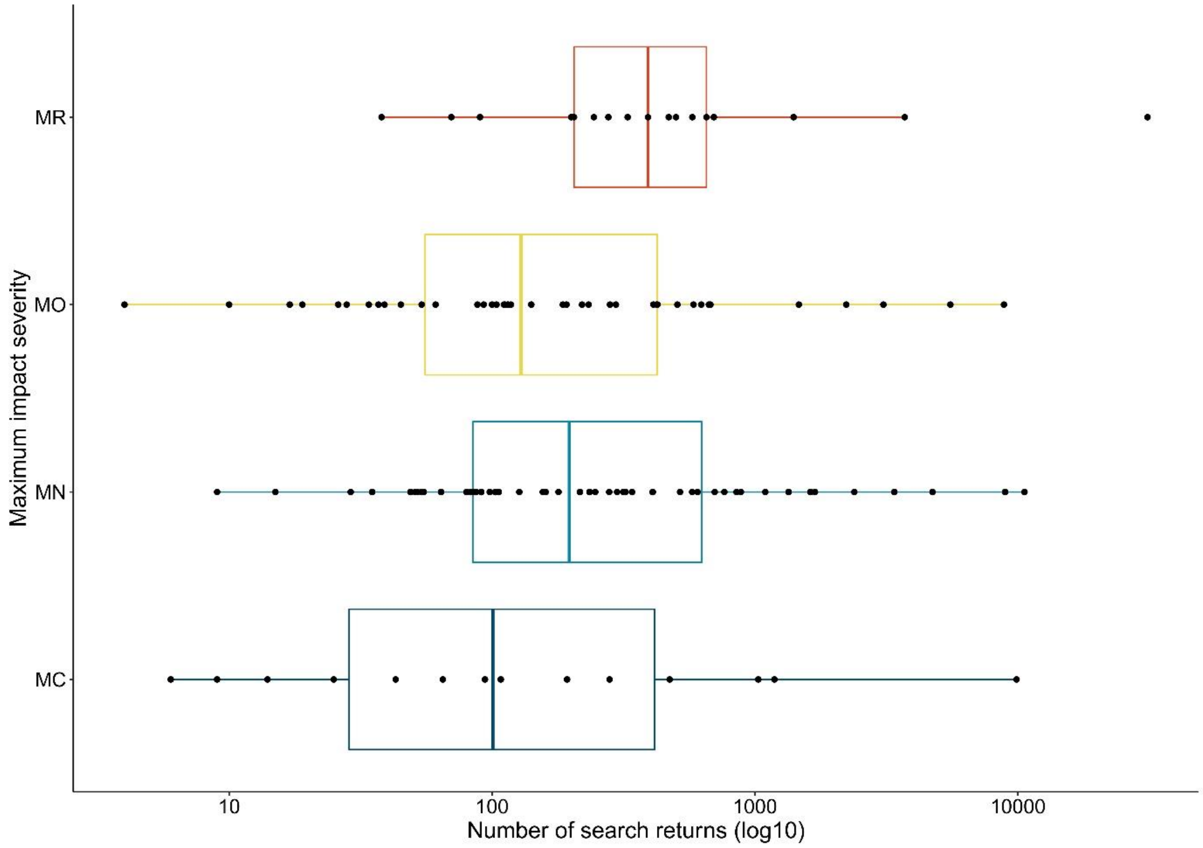


Figure S4. Relationship between the number of Web of Science literature search returns (log10) for each alien insect species with evidence of environmental impact and their global maximum impact severity.

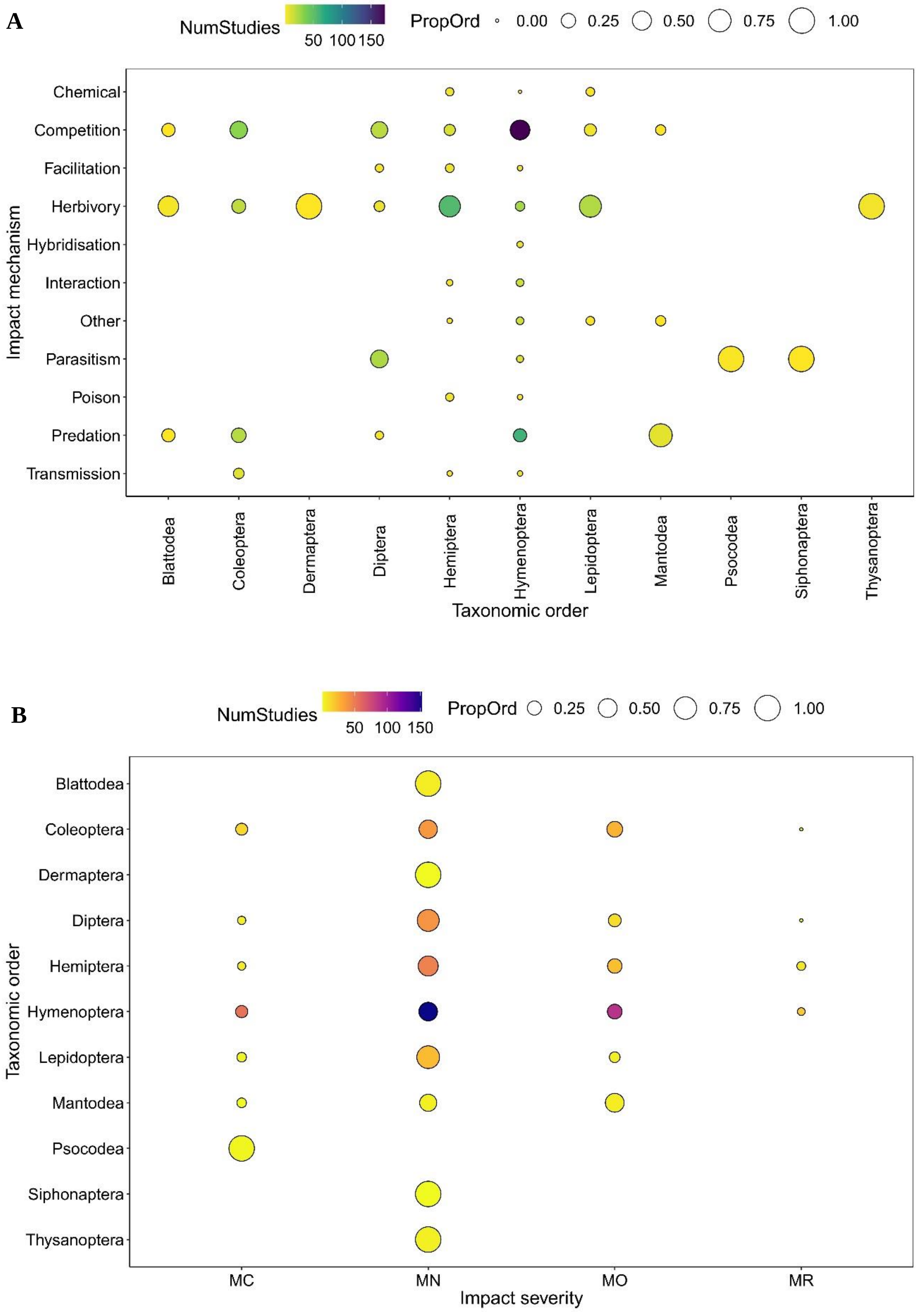


Figure S3. Taxonomic distribution of the mechanisms and severities of environmental impacts of alien insects. A. Proportion of impact mechanism attribution within taxonomic order, i.e., the proportion of an impact mechanism attributed to a given taxonomic order within a given taxonomic order. Colour represents the number of individual studies, and circle size represents the proportion of individual studies, where a given mechanism was attributed. For example, environmental impact evidence for Dermaptera, Psocodea, Siphonaptera, and Thysanoptera was represented by a single impact mechanism, hence the large circle size. Impact mechanisms are Chemical/Physical/Structural impact on ecosystem, Competition, Facilitation of native species, Herbivory, Hybridization, Interaction with other aliens, Other, Parasitism, Poisoning/toxicity, Predation, and Transmission of disease. B. Proportion of impact severity attribution within taxonomic order, i.e., the proportion of a given impact severity attributed to a given taxonomic order across all taxonomic orders. Colour represents the number of individual studies, and size represents the proportion of individual studies, where a given severity was attributed. For example, with the exception of Coleoptera and Mantodea, environmental impact evidence for most taxonomic orders was dominated by a single impact severity Impact severity categories are Minimal Concern (MC, impacts on native taxa negligible), Minor (MN, no evidence for a decline in population sizes of native taxa), Moderate (MO, impact native species population sizes but no evidence of local apparent extinction), Major (MR, reversible local extinction of one or more native taxa) (there were no species assessed as having a Massive (MV) impact, irreversible local extinction of one or more native taxa) (for full descriptions of categories used see Clarke et al. 2021). Note: proportions of 0.00 indicate a small non-zero number. True zeros are indicated by the absence of a circle on the plot.

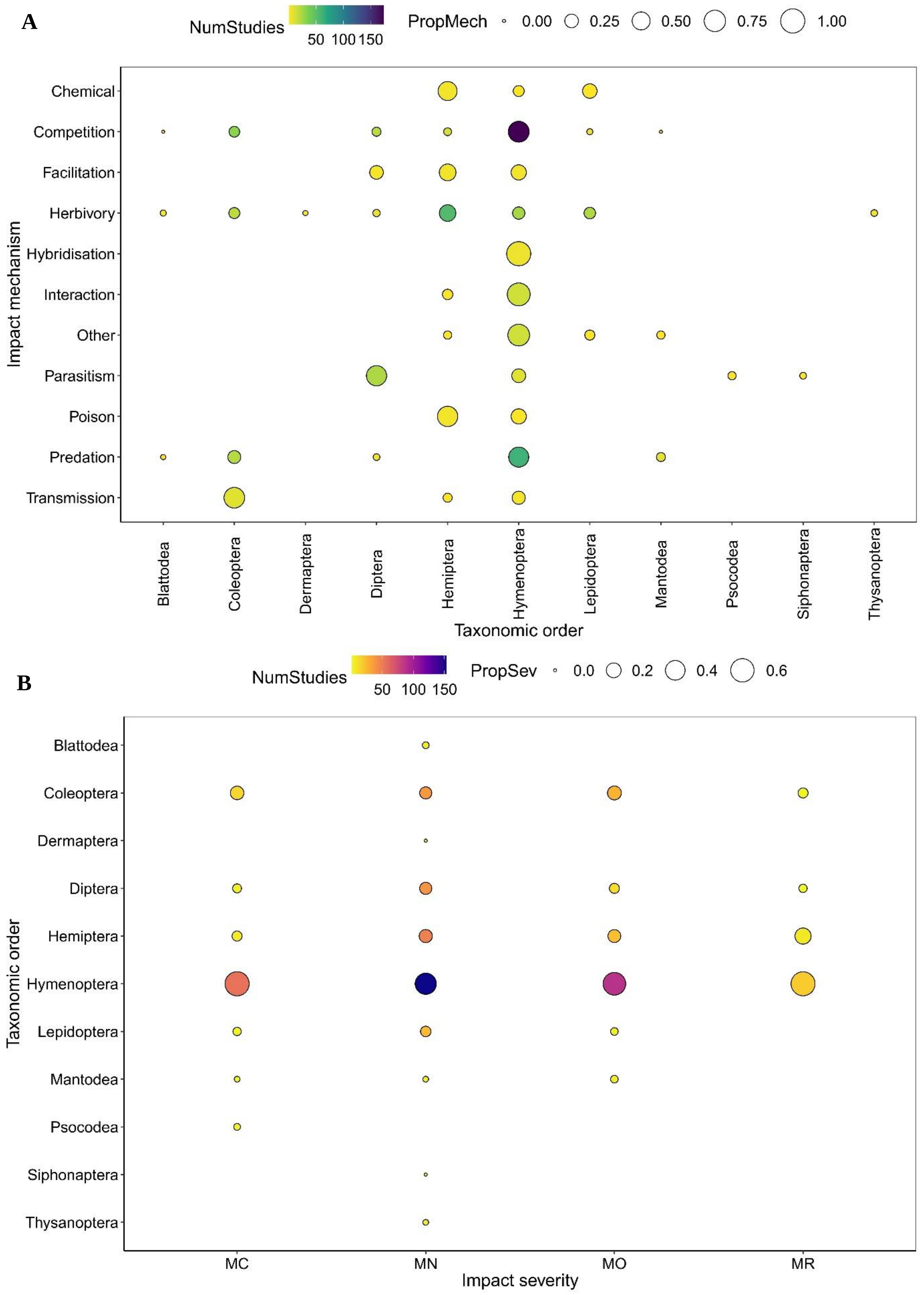


Figure S4. Taxonomic distribution of the mechanisms and severities of environmental impacts of alien insects. A. Proportion of impact mechanism attribution among taxonomic order, i.e., the proportion of an impact mechanism attributed to a given taxonomic order across all taxonomic orders. Colour represents the number of individual studies, and circle size represents the proportion of individual studies, where a given mechanism was attributed. For example, Hymenoptera was the only order with species impacting the environment via Hybridization, hence the large circle size. Coleoptera had the largest proportion of impacts via Transmission of disease. Impact mechanisms are Chemical/Physical/Structural impact on ecosystem, Competition, Facilitation of native species, Herbivory, Hybridization, Interaction with other aliens, Other, Parasitism, Poisoning/toxicity, Predation, and Transmission of disease. B. Proportion of impact severity attribution among taxonomic order, i.e., the proportion of a given impact severity attributed to a given taxonomic order across all taxonomic orders. Colour represents the number of individual studies, and size represents the proportion of individual studies, where a given severity was attributed. For example, the largest proportion of impact evidence for each impact severity category related to insects from Hymenoptera. Impact severity categories are Minimal Concern (MC, impacts on native taxa negligible), Minor (MN, no evidence for a decline in population sizes of native taxa), Moderate (MO, impact native species population sizes but no evidence of local apparent extinction), Major (MR, reversible local extinction of one or more native taxa) (there were no species assessed as having a Massive (MV) impact, irreversible local extinction of one or more native taxa) (for full descriptions of categories used see Clarke et al. 2021). Note: proportions of 0.00 indicate a small non-zero number. True zeros are indicated by the absence of a circle on the plot.

Table S1. Global maximum impact severity and associated impact mechanism for each insect species with known or presumed alien populations. Impact severity: Minimal Concern (MC), Minor (MN), Moderate (MO), Major (MR), Data Deficient (DD).

| **Order** | **Family** | **Scientific name** | **Mechanism** | **Severity** |
|---|---|---|---|---|
| Coleoptera | Carabidae | *Merizodus soledadinus* | Predation | MR |
| Coleoptera | Scarabaeidae | *Digitonthophagus gazella* | Competition | MR |
| Diptera | Calliphoridae | *Chrysomya albiceps* | Competition | MR |
| Hemiptera | Diaspididae | *Aulacaspis yasumatsui* | Herbivory | MR |
| Hemiptera | Margarodidae | *Icerya purchasi* | Herbivory | MR |
| Hymenoptera | Formicidae | *Anoplolepis gracilipes* | Competition | MR |
| Hymenoptera | Formicidae | *Myrmica rubra* | Competition | MR |
| Coleoptera | Carabidae | *Cicindela trifasciata* | Competition | MO |
| Coleoptera | Carabidae | *Trechisibus antarcticus* | Predation | MO |
| Coleoptera | Carabidae | *Trechus obtusus* | Competition | MO |
| Coleoptera | Cerambycidae | *Tetropium fuscum* | Competition | MO |
| Coleoptera | Coccinellidae | *Harmonia axyridis* | Competition | MO |
| Coleoptera | Curculionidae | *Larinus planus* | Herbivory | MO |
| Coleoptera | Curculionidae | *Xyleborus glabratus* | Transmission of disease | MO |
| Diptera | Culicidae | *Ochlerotatus japonicus* | Competition | MO |
| Diptera | Tachinidae | *Bessa remota* | Predation | MO |
| Diptera | Tachinidae | *Trichopoda pilipes* | Parasitism | MO |
| Hemiptera | Adelgidae | *Pineus pini* | Competition | MO |
| Hemiptera | Aleyrodidae | *Bemisia tabaci* | Competition | MO |
| Hemiptera | Coccidae | *Toumeyella parvicornis* | Herbivory | MO |
| Hemiptera | Corixidae | *Trichocorixa verticalis verticalis* | Competition | MO |
| Hemiptera | Ortheziidae | *Insignorthezia insignis* | Herbivory | MO |
| Hymenoptera | Aphelinidae | *Cales noacki* | Competition | MO |
| Hymenoptera | Apidae | *Bombus ruderatus* | Competition | MO |
| Hymenoptera | Apidae | *Bombus terrestris* | Competition | MO |
| Hymenoptera | Braconidae | *Lysiphlebus testaceipes* | Competition | MO |
| Hymenoptera | Diprionidae | *Gilpinia hercyniae* | Herbivory | MO |
| Hymenoptera | Encyrtidae | *Metaphycus helvolus* | Competition | MO |
| Hymenoptera | Eulophidae | *Citrostichus phyllocnistoides* | Competition | MO |
| Hymenoptera | Eulophidae | *Quadrastichus erythrinae* | Parasitism | MO |
| Hymenoptera | Eumenidae | *Polistes dominula* | Competition | MO |
| Hymenoptera | Formicidae | *Lasius neglectus* | Competition | MO |

| | | | | |
|---|---|---|---|---|
| Hymenoptera | Formicidae | *Pachycondyla chinensis* | Competition | MO |
| Hymenoptera | Formicidae | *Paratrechina fulva* | Competition | MO |
| Hymenoptera | Formicidae | *Tapinoma melanocephalum* | Predation | MO |
| Hymenoptera | Formicidae | *Tetramorium bicarinatum* | Interaction with other alien species | MO |
| Hymenoptera | Formicidae | *Wasmannia auropunctata* | Competition | MO |
| Hymenoptera | Siricidae | *Sirex noctilio* | Herbivory | MO |
| Hymenoptera | Torymidae | *Torymus sinensis* | Hybridisation | MO |
| Lepidoptera | Crambidae | *Chilo partellus* | Competition | MO |
| Blattodea | Blattidae | *Periplaneta americana* | Predation | MN |
| Blattodea | Blattidae | *Periplaneta australasiae* | Predation | MN |
| Blattodea | Rhinotermitidae | *Coptotermes formosanus* | Herbivory | MN |
| Blattodea | Rhinotermitidae | *Coptotermes gestroi* | Herbivory | MN |
| Blattodea | Rhinotermitidae | *Reticulitermes flavipes* | Competition | MN |
| Coleoptera | Carabidae | *Pterostichus melanarius* | Competition | MN |
| Coleoptera | Cerambycidae | *Anoplophora glabripennis* | Herbivory | MN |
| Coleoptera | Curculionidae | *Rhinocyllus conicus* | Herbivory | MN |
| Coleoptera | Curculionidae | *Xylosandrus compactus* | Herbivory | MN |
| Coleoptera | Dryophthoridae | *Scyphophorus acupunctatus* | Herbivory | MN |
| Coleoptera | Nitidulidae | *Aethina tumida* | Predation | MN |
| Dermaptera | Forficulidae | *Forficula auricularia* | Herbivory | MN |
| Diptera | Calliphoridae | *Calliphora vicina* | Competition | MN |
| Diptera | Cecidomyiidae | *Contarinia baeri* | Herbivory | MN |
| Diptera | Cecidomyiidae | *Obolodiplosis robiniae* | Herbivory | MN |
| Diptera | Culicidae | *Aedes albopictus* | Competition | MN |
| Diptera | Muscidae | *Philornis downsi* | Parasitism | MN |
| Diptera | Tephritidae | *Bactrocera cucurbitae* | Competition | MN |
| Diptera | Tephritidae | *Urophora affinis* | Facilitation of native species | MN |
| Diptera | Tephritidae | *Urophora quadrifasciata* | Facilitation of native species | MN |
| Hemiptera | Adelgidae | *Adelges tsugae* | Herbivory | MN |
| Hemiptera | Aphididae | *Aphis gossypii* | Herbivory | MN |
| Hemiptera | Aphididae | *Aphis spiraecola* | Herbivory | MN |
| Hemiptera | Aphididae | *Chaetosiphon fragaefolii* | Transmission of disease | MN |
| Hemiptera | Aphididae | *Cinara cupressi* | Herbivory | MN |
| Hemiptera | Aphididae | *Myzus ascalonicus* | Herbivory | MN |
| Hemiptera | Aphididae | *Uroleucon nigrotuberculatum* | Facilitation of native species | MN |
| Hemiptera | Pseudococcidae | *Phenacoccus solenopsis* | Facilitation of native species | MN |

| | | | | |
|---|---|---|---|---|
| Hymenoptera | Apidae | *Apis cerana* | Competition | MN |
| Hymenoptera | Apidae | *Apis mellifera* | Interaction with other alien species | MN |
| Hymenoptera | Apidae | *Apis mellifera scutellata* | Competition | MN |
| Hymenoptera | Apidae | *Bombus impatiens* | Competition | MN |
| Hymenoptera | Apidae | *Bombus lucorum* | Competition | MN |
| Hymenoptera | Argidae | *Aproceros leucopoda* | Herbivory | MN |
| Hymenoptera | Braconidae | *Microctonus aethiopoides* | Parasitism | MN |
| Hymenoptera | Cynipidae | *Andricus quercuscalicis* | Other | MN |
| Hymenoptera | Cynipidae | *Dryocosmus kuriphilus* | Herbivory | MN |
| Hymenoptera | Eumenidae | *Polistes chinensis antennalis* | Predation | MN |
| Hymenoptera | Formicidae | *Linepithema humile* | Chemical/physical/structural impact on ecosystem | MN |
| Hymenoptera | Formicidae | *Monomorium destructor* | Competition | MN |
| Hymenoptera | Formicidae | *Monomorium pharaonis* | Predation | MN |
| Hymenoptera | Formicidae | *Paratrechina longicornis* | Competition | MN |
| Hymenoptera | Formicidae | *Paratrechina pubens* | Competition | MN |
| Hymenoptera | Formicidae | *Pheidole megacephala* | Predation | MN |
| Hymenoptera | Formicidae | *Solenopsis geminata* | Predation | MN |
| Hymenoptera | Formicidae | *Solenopsis invicta* | Facilitation of native species | MN |
| Hymenoptera | Formicidae | *Solenopsis papuana* | Competition | MN |
| Hymenoptera | Formicidae | *Technomyrmex albipes* | Facilitation of native species | MN |
| Hymenoptera | Megachilidae | *Megachile apicalis* | Competition | MN |
| Hymenoptera | Megachilidae | *Megachile rotundata* | Competition | MN |
| Hymenoptera | Pamphiliidae | *Acantholyda erythrocephala* | Herbivory | MN |
| Hymenoptera | Pteromalidae | *Pteromalus puparum* | Parasitism | MN |
| Hymenoptera | Scelionidae | *Trissolcus basalis* | Parasitism | MN |
| Hymenoptera | Tenthredinidae | *Fenusa pusilla* | Herbivory | MN |
| Hymenoptera | Vespidae | *Vespa velutina* | Competition | MN |
| Hymenoptera | Vespidae | *Vespula pensylvanica* | Transmission of disease | MN |
| Hymenoptera | Vespidae | *Vespula vulgaris* | Other | MN |
| Lepidoptera | Coleophoridae | *Coleophora laricella* | Herbivory | MN |
| Lepidoptera | Crambidae | *Cydalima perspectalis* | Herbivory | MN |
| Lepidoptera | Erebidae | *Hyphantria cunea* | Herbivory | MN |
| Lepidoptera | Erebidae | *Lymantria dispar* | Herbivory | MN |
| Lepidoptera | Geometridae | *Operophtera brumata* | Herbivory | MN |
| Lepidoptera | Pyralidae | *Cactoblastis cactorum* | Herbivory | MN |
| Mantodea | Mantidae | *Tenodera sinensis* | Predation | MN |
| Siphonaptera | Leptopsyllidae | *Leptopsylla segnis* | Parasitism | MN |

| | | | | |
|---|---|---|---|---|
| Thysanoptera | Thripidae | *Taeniothrips inconsequens* | Herbivory | MN |
| Coleoptera | Brachyceridae | *Stenopelmus rufinasus* | Herbivory | MC |
| Coleoptera | Cerambycidae | *Phoracantha semipunctata* | Herbivory | MC |
| Diptera | Drosophilidae | *Drosophila subobscura* | Competition | MC |
| Hemiptera | Adelgidae | *Adelges abietis* | Herbivory | MC |
| Hemiptera | Adelgidae | *Adelges cooleyi* | Herbivory | MC |
| Hymenoptera | Apidae | *Apis mellifera carnica* | Hybridisation | MC |
| Hymenoptera | Apidae | *Bombus hortorum* | Interaction with other alien species | MC |
| Hymenoptera | Apidae | *Bombus pascuorum* | Competition | MC |
| Hymenoptera | Cynipidae | *Andricus corruptrix* | Competition | MC |
| Hymenoptera | Cynipidae | *Andricus kollari* | Competition | MC |
| Hymenoptera | Cynipidae | *Andricus lignicolus* | Competition | MC |
| Hymenoptera | Diprionidae | *Diprion similis* | Transmission of disease | MC |
| Hymenoptera | Formicidae | *Monomorium floricola* | Competition | MC |
| Hymenoptera | Megachilidae | *Afranthidium repetitum* | Competition | MC |
| Hymenoptera | Tenthredinidae | *Profenusa thomsoni* | Herbivory | MC |
| Hymenoptera | Vespidae | *Vespula germanica* | Competition | MC |
| Lepidoptera | Noctuidae | *Helicoverpa armigera* | Competition | MC |
| Mantodea | Mantidae | *Mantis religiosa* | Predation | MC |
| Psocodea | Gyropidae | *Pitrufquenia coypus* | Parasitism | MC |
| Blattodea | Blaberidae | *Pycnoscelus surinamensis* | NA | DD |
| Blattodea | Ectobiidae | *Blattella germanica* | NA | DD |
| Blattodea | Ectobiidae | *Supella longipalpa* | NA | DD |
| Blattodea | Kalotermitidae | *Cryptotermes brevis* | NA | DD |
| Coleoptera | Anobiidae | *Stegobium paniceum* | NA | DD |
| Coleoptera | Bostrichidae | *Bostrichus cembrae* | NA | DD |
| Coleoptera | Bostrichidae | *Rhyzopertha dominica* | NA | DD |
| Coleoptera | Buprestidae | *Agrilus ribesi* | NA | DD |
| Coleoptera | Buprestidae | *Agrilus sinuatus* | NA | DD |
| Coleoptera | Cerambycidae | *Acrocinus longimanus* | NA | DD |
| Coleoptera | Cerambycidae | *Anoplophora chinensis* | NA | DD |
| Coleoptera | Cerambycidae | *Monochamus sutor* | NA | DD |
| Coleoptera | Chrysomelidae | *Acanthoscelides obtectus* | NA | DD |
| Coleoptera | Chrysomelidae | *Bruchus pisorum* | NA | DD |
| Coleoptera | Chrysomelidae | *Bruchus rufimanus* | NA | DD |
| Coleoptera | Chrysomelidae | *Callosobruchus chinensis* | NA | DD |
| Coleoptera | Chrysomelidae | *Leptinotarsa decemlineata* | NA | DD |
| Coleoptera | Coccinellidae | *Propylea quatuordecimpunctata* | NA | DD |

| | | | | |
|---|---|---|---|---|
| Coleoptera | Curculionidae | *Anthonomus grandis* | NA | DD |
| Coleoptera | Curculionidae | *Hylurgus rufipennis* | NA | DD |
| Coleoptera | Curculionidae | *Orthotomicus erosus* | NA | DD |
| Coleoptera | Curculionidae | *Otiorhynchus ovatus* | NA | DD |
| Coleoptera | Curculionidae | *Xylosandrus morigerus* | NA | DD |
| Coleoptera | Dermestidae | *Thylodrias contractus* | NA | DD |
| Coleoptera | Dermestidae | *Trogoderma granarium* | NA | DD |
| Coleoptera | Dryophthoridae | *Sitophilus oryzae* | NA | DD |
| Coleoptera | Dynastidae | *Heteronychus arator* | NA | DD |
| Coleoptera | Dytiscidae | *Cybister lateralimarginalis* | NA | DD |
| Coleoptera | Laemophloeidae | *Cryptolestes pusillus* | NA | DD |
| Coleoptera | Laemophloeidae | *Cryptolestes turcicus* | NA | DD |
| Coleoptera | Lucanidae | *Prosopocoilus inclinatus* | NA | DD |
| Coleoptera | Nitidulidae | *Stelidota geminata* | NA | DD |
| Coleoptera | Rutelidae | *Adoretus versutus* | NA | DD |
| Coleoptera | Scarabaeidae | *Onthophagus vacca* | NA | DD |
| Coleoptera | Tenebrionidae | *Tribolium castaneum* | NA | DD |
| Coleoptera | Tenebrionidae | *Tribolium destructor* | NA | DD |
| Dermaptera | Anisolabididae | *Anisolabis maritima* | NA | DD |
| Dermaptera | Anisolabididae | *Euborellia annulipes* | NA | DD |
| Dermaptera | Anisolabididae | *Gonolabis marginalis* | NA | DD |
| Dermaptera | Chelisochidae | *Chelisoches morio* | NA | DD |
| Dermaptera | Forficulidae | *Doru taeniatum* | NA | DD |
| Dermaptera | Forficulidae | *Forficula decipiens* | NA | DD |
| Dermaptera | Labiduridae | *Labidura riparia* | NA | DD |
| Dermaptera | Labiduridae | *Nala lividipes* | NA | DD |
| Dermaptera | Spongiphoridae | *Labia minor* | NA | DD |
| Dermaptera | Spongiphoridae | *Marava arachidis* | NA | DD |
| Diptera | Agromyzidae | *Liriomyza huidobrensis* | NA | DD |
| Diptera | Agromyzidae | *Liriomyza sativae* | NA | DD |
| Diptera | Agromyzidae | *Liriomyza trifolii* | NA | DD |
| Diptera | Agromyzidae | *Paraphytomyza populicola* | NA | DD |
| Diptera | Calliphoridae | *Calliphora vomitoria* | NA | DD |
| Diptera | Calliphoridae | *Chrysomya chloropyga* | NA | DD |
| Diptera | Calliphoridae | *Chrysomya megacephala* | NA | DD |
| Diptera | Calliphoridae | *Lucilia sericata* | NA | DD |
| Diptera | Cecidomyiidae | *Contarinia citri* | NA | DD |
| Diptera | Ceratopogonidae | *Culicoides kingi* | NA | DD |

| | | | | |
|---|---|---|---|---|
| Diptera | Drosophilidae | *Scaptodrosophila latifasciaeformis* | NA | DD |
| Diptera | Drosophilidae | *Zaprionus indianus* | NA | DD |
| Diptera | Lauxaniidae | *Meiosimyza rorida* | NA | DD |
| Diptera | Muscidae | *Hydrotaea aenescens* | NA | DD |
| Diptera | Syrphidae | *Eristalis intricaria* | NA | DD |
| Diptera | Tachinidae | *Nealsomyia rufella* | NA | DD |
| Diptera | Tephritidae | *Anastrepha obliqua* | NA | DD |
| Diptera | Tephritidae | *Bactrocera tryoni* | NA | DD |
| Diptera | Tephritidae | *Ceratitis capitata* | NA | DD |
| Diptera | Tephritidae | *Ceratitis rosa* | NA | DD |
| Embioptera | Embiidae | *Parembia persica* | NA | DD |
| Embioptera | Oligotomidae | *Haploembia solieri* | NA | DD |
| Embioptera | Oligotomidae | *Oligotoma michaeli* | NA | DD |
| Embioptera | Oligotomidae | *Oligotoma nigra* | NA | DD |
| Embioptera | Oligotomidae | *Oligotoma saundersii* | NA | DD |
| Ephemeroptera | Baetidae | *Baetis liebenauae* | NA | DD |
| Ephemeroptera | Baetidae | *Baetis tracheatus* | NA | DD |
| Hemiptera | Aleyrodidae | *Acaudaleyrodes rachipora* | NA | DD |
| Hemiptera | Aleyrodidae | *Aleurodicus floccissimus* | NA | DD |
| Hemiptera | Aleyrodidae | *Aleurothrixus floccosus* | NA | DD |
| Hemiptera | Aleyrodidae | *Aleurotrachelus atratus* | NA | DD |
| Hemiptera | Aleyrodidae | *Trialeurodes vaporariorum* | NA | DD |
| Hemiptera | Aphalaridae | *Ctenarytaina eucalypti* | NA | DD |
| Hemiptera | Aphididae | *Aphis forbesi* | NA | DD |
| Hemiptera | Aphididae | *Chromaphis juglandicola* | NA | DD |
| Hemiptera | Aphididae | *Macrosiphoniella sanborni* | NA | DD |
| Hemiptera | Aphididae | *Myzus varians* | NA | DD |
| Hemiptera | Aphididae | *Panaphis juglandis* | NA | DD |
| Hemiptera | Aphididae | *Reticulaphis distylii* | NA | DD |
| Hemiptera | Aphididae | *Rhopalosiphum maidis* | NA | DD |
| Hemiptera | Aphididae | *Sarucallis kahawaluokalani* | NA | DD |
| Hemiptera | Cicadellidae | *Empoasca fabalis* | NA | DD |
| Hemiptera | Cicadellidae | *Erasmoneura variabilis* | NA | DD |
| Hemiptera | Cicadellidae | *Homalodisca vitripennis* | NA | DD |
| Hemiptera | Coccidae | *Pulvinaria regalis* | NA | DD |
| Hemiptera | Coccidae | *Saissetia oleae* | NA | DD |
| Hemiptera | Diaspididae | *Aspidiotus nerii* | NA | DD |

| | | | | |
|---|---|---|---|---|
| Hemiptera | Diaspididae | *Hemiberlesia pitysophila* | NA | DD |
| Hemiptera | Liviidae | *Diaphorina citri* | NA | DD |
| Hemiptera | Membracidae | *Stictocephala bisonia* | NA | DD |
| Hemiptera | Pentatomidae | *Halyomorpha halys* | NA | DD |
| Hemiptera | Pseudococcidae | *Balanococcus diminutus* | NA | DD |
| Hemiptera | Pseudococcidae | *Oracella acuta* | NA | DD |
| Hemiptera | Pseudococcidae | *Pseudococcus calceolariae* | NA | DD |
| Hemiptera | Pseudococcidae | *Pseudococcus viburni* | NA | DD |
| Hemiptera | Pseudococcidae | *Tipula paludosa* | NA | DD |
| Hemiptera | Psyllidae | *Cacopsylla fulguralis* | NA | DD |
| Hemiptera | Psyllidae | *Heteropsylla cubana* | NA | DD |
| Hemiptera | Reduviidae | *Agriosphodrus dohrni* | NA | DD |
| Hemiptera | Tingidae | *Corythucha ciliata* | NA | DD |
| Hymenoptera | Aphelinidae | *Aphytis mytilaspidis* | NA | DD |
| Hymenoptera | Aphelinidae | *Encarsia guadeloupae* | NA | DD |
| Hymenoptera | Apidae | *Bombus hypnorum* | NA | DD |
| Hymenoptera | Argidae | *Arge berberidis* | NA | DD |
| Hymenoptera | Diprionidae | *Neodiprion sertifer* | NA | DD |
| Hymenoptera | Eulophidae | *Cirrospilus ingenuus* | NA | DD |
| Hymenoptera | Eulophidae | *Leptocybe invasa* | NA | DD |
| Hymenoptera | Eulophidae | *Ophelimus maskelli* | NA | DD |
| Hymenoptera | Eulophidae | *Semielacher petiolata* | NA | DD |
| Hymenoptera | Eulophidae | *Thripobius javae* | NA | DD |
| Hymenoptera | Eumenidae | *Delta pyriforme* | NA | DD |
| Hymenoptera | Eumenidae | *Polistes olivaceus* | NA | DD |
| Hymenoptera | Formicidae | *Acromyrmex octospinosus* | NA | DD |
| Hymenoptera | Formicidae | *Cardiocondyla emeryi* | NA | DD |
| Hymenoptera | Formicidae | *Cardiocondyla obscurior* | NA | DD |
| Hymenoptera | Formicidae | *Hypoponera punctatissima* | NA | DD |
| Hymenoptera | Formicidae | *Iridomyrmex anceps* | NA | DD |
| Hymenoptera | Formicidae | *Lasius niger* | NA | DD |
| Hymenoptera | Formicidae | *Lepisiota frauenfeldi* | NA | DD |
| Hymenoptera | Formicidae | *Monomorium indicum* | NA | DD |
| Hymenoptera | Formicidae | *Pachycondyla sennaarensis* | NA | DD |
| Hymenoptera | Formicidae | *Paratrechina flavipes* | NA | DD |
| Hymenoptera | Formicidae | *Paratrechina jaegerskioeldi* | NA | DD |
| Hymenoptera | Formicidae | *Pheidole teneriffana* | NA | DD |
| Hymenoptera | Formicidae | *Solenopsis richteri* | NA | DD |
| Hymenoptera | Formicidae | *Tapinoma simrothi* | NA | DD |

| | | | | |
|---|---|---|---|---|
| Hymenoptera | Formicidae | *Tetramorium caldarium* | NA | DD |
| Hymenoptera | Pamphiliidae | *Cephalcia lariciphila* | NA | DD |
| Hymenoptera | Scelionidae | *Sceliphron caementarium* | NA | DD |
| Hymenoptera | Scelionidae | *Sceliphron curvatum* | NA | DD |
| Hymenoptera | Scelionidae | *Sceliphron deforme* | NA | DD |
| Hymenoptera | Siricidae | *Sirex cyaneus* | NA | DD |
| Hymenoptera | Tenthredinidae | *Nematus ribesii* | NA | DD |
| Hymenoptera | Tenthredinidae | *Pristiphora erichsonii* | NA | DD |
| Hymenoptera | Tenthredinidae | *Pristiphora geniculata* | NA | DD |
| Hymenoptera | Vespidae | *Dolichovespula norwegica* | NA | DD |
| Hymenoptera | Vespidae | *Vespa mandarinia* | NA | DD |
| Hymenoptera | Vespidae | *Vespa velutina nigrithorax* | NA | DD |
| Hymenoptera | Vespidae | *Vespula rufa* | NA | DD |
| Lepidoptera | Cosmopterigidae | *Anatrachyntis badia* | NA | DD |
| Lepidoptera | Crambidae | *Spoladea recurvalis* | NA | DD |
| Lepidoptera | Erebidae | *Lymantria mathura* | NA | DD |
| Lepidoptera | Gelechiidae | *Pectinophora gossypiella* | NA | DD |
| Lepidoptera | Gelechiidae | *Sitotroga cerealella* | NA | DD |
| Lepidoptera | Gelechiidae | *Tuta absoluta* | NA | DD |
| Lepidoptera | Geometridae | *Bupalus piniaria* | NA | DD |
| Lepidoptera | Gracillariidae | *Phyllocnistis citrella* | NA | DD |
| Lepidoptera | Limacodidae | *Parasa lepida* | NA | DD |
| Lepidoptera | Lycaenidae | *Virachola livia* | NA | DD |
| Lepidoptera | Lycaenidae | *Zizina labradus* | NA | DD |
| Lepidoptera | Noctuidae | *Penicillaria jocosatrix* | NA | DD |
| Lepidoptera | Noctuidae | *Spodoptera frugiperda* | NA | DD |
| Lepidoptera | Notodontidae | *Thaumetopoea processionea* | NA | DD |
| Lepidoptera | Pieridae | *Pieris rapae* | NA | DD |
| Lepidoptera | Plutellidae | *Plutella xylostella* | NA | DD |
| Lepidoptera | Pyralidae | *Cadra cautella* | NA | DD |
| Lepidoptera | Pyralidae | *Cadra figulilella* | NA | DD |
| Lepidoptera | Tortricidae | *Grapholita molesta* | NA | DD |
| Neuroptera | Chrysopidae | *Chrysoperla rufilabris* | NA | DD |
| Neuroptera | Hemerobiidae | *Hemerobius bolivari* | NA | DD |
| Odonata | Coenagrionidae | *Pseudagrion microcephalum* | NA | DD |
| Orthoptera | Acrididae | *Locusta migratoria* | NA | DD |
| Orthoptera | Gryllidae | *Acheta domesticus* | NA | DD |
| Phasmatodea | Bacillidae | *Bacillus rossius* | NA | DD |
| Phasmatodea | Bacillidae | *Clonopsis gallica* | NA | DD |

| | | | | |
|---|---|---|---|---|
| Phasmatodea | Diapheromeridae | *Neohirasea japonica* | NA | DD |
| Phasmatodea | Phasmatidae | *Acanthoxyla inermis* | NA | DD |
| Phasmatodea | Phasmatidae | *Carausius morosus* | NA | DD |
| Phasmatodea | Phasmatidae | *Clitarchus hookeri* | NA | DD |
| Psocodea | Ectopsocidae | *Ectopsocus briggsi* | NA | DD |
| Psocodea | Gyropidae | *Gyropus ovalis* | NA | DD |
| Psocodea | Lachesillidae | *Lachesilla greeni* | NA | DD |
| Psocodea | Lachesillidae | *Lachesilla tectorum* | NA | DD |
| Psocodea | Lepidopsocidae | *Echmepteryx madagascariensis* | NA | DD |
| Psocodea | Liposcelididae | *Liposcelis corrodens* | NA | DD |
| Psocodea | Liposcelididae | *Liposcelis decolor* | NA | DD |
| Psocodea | Liposcelididae | *Liposcelis kidderi* | NA | DD |
| Psocodea | Philopteridae | *Cuclotogaster heterographus* | NA | DD |
| Psocodea | Philopteridae | *Goniocotes rectangulatus* | NA | DD |
| Psocodea | Philopteridae | *Goniodes gigas* | NA | DD |
| Psocodea | Philopteridae | *Lipeurus caponis* | NA | DD |
| Psocodea | Philopteridae | *Lipeurus maculosus* | NA | DD |
| Psocodea | Psyllipsocidae | *Dorypteryx domestica* | NA | DD |
| Psocodea | Sphaeropsocidae | *Badonnelia titei* | NA | DD |
| Psocodea | Trichodectidae | *Stachiella octomaculatus* | NA | DD |
| Psocodea | Trogiidae | *Lepinotus patruelis* | NA | DD |
| Psocodea | Trogiidae | *Trogium pulsatorium* | NA | DD |
| Siphonaptera | Ceratophyllidae | *Nosopsyllus fasciatus* | NA | DD |
| Siphonaptera | Ceratophyllidae | *Nosopsyllus londinensis* | NA | DD |
| Siphonaptera | Pulicidae | *Xenopsylla cheopis* | NA | DD |
| Thysanoptera | Phlaeothripidae | *Gynaikothrips ficorum* | NA | DD |
| Thysanoptera | Thripidae | *Chaetanaphothrips orchidii* | NA | DD |
| Thysanoptera | Thripidae | *Frankliniella fusca* | NA | DD |
| Thysanoptera | Thripidae | *Frankliniella schultzei* | NA | DD |
| Thysanoptera | Thripidae | *Microcephalothrips abdominalis* | NA | DD |
| Thysanoptera | Thripidae | *Parthenothrips dracaenae* | NA | DD |
| Thysanoptera | Thripidae | *Pezothrips kellyanus* | NA | DD |
| Thysanoptera | Thripidae | *Thrips palmi* | NA | DD |
| Thysanoptera | Thripidae | *Thrips simplex* | NA | DD |
| Zygentoma | Lepismatidae | *Ctenolepisma lineata* | NA | DD |
| Zygentoma | Lepismatidae | *Lepisma saccharina* | NA | DD |
| Zygentoma | Lepismatidae | *Thermobia domestica* | NA | DD |
| Zygentoma | Nicoletiidae | *Coletinia maggii* | NA | DD |

| | | | | |
|---|---|---|---|---|
| Zygentoma | Nicoletiidae | *Nicoletia phytophila* | NA | DD |
| Zygentoma | Nicoletiidae | *Proatelurina pseudolepisma* | NA | DD |
| Coleoptera | Chrysomelidae | *Phratora vitellinae* | NA | NA |
| Dermaptera | Anisolabididae | *Carcinophora americana* | NA | NA |
| Dermaptera | Anisolabididae | *Euborellia janeirensis* | NA | NA |
| Dermaptera | Anisolabididae | *Euborellia peregrina* | NA | NA |
| Dermaptera | Forficulidae | *Kleter devians* | NA | NA |
| Dermaptera | Forficulidae | *Metresura ruficeps* | NA | NA |
| Dermaptera | Labiduridae | *Forcipula gariazzi* | NA | NA |
| Diptera | Culicidae | *Anopheles quadrimaculatus* | NA | NA |
| Lepidoptera | Cossidae | *Coryphodema tristis* | NA | NA |
| Lepidoptera | Erebidae | *Orgyia thyellina* | NA | NA |
| Lepidoptera | Lasiocampidae | *Dendrolimus superans sibiricus* | NA | NA |
| Lepidoptera | Saturniidae | *Ormiscodes amphimone* | NA | NA |
| Psocodea | Caeciliusidae | *Valenzuela burmeisteri* | NA | NA |
| Trichoptera | Hydropsychidae | *Hydropsyche bulgaromanorum* | NA | NA |
| Trichoptera | Hydropsychidae | *Hydropsyche contubernalis* | NA | NA |
| Trichoptera | Leptoceridae | *Leptocerus lusitanicus* | NA | NA |

Table S2. Intraspecific environmental impact information for the seven most studied alien insect species in this assessment. The range of values demonstrate the variability and depth of the collected information.

| Species | Common name | Number publications | Mechanisms | Severities | Confidence | Max severity |
|---|---|---|---|---|---|---|
| *Harmonia axyridis* | Harlequin lady beetle | 33 | Competition, Predation | Minimal Concern, Minor, Moderate | Low, Medium, High | Moderate |
| *Philornis downsi* | NA | 18 | Parasitism | Minor, Moderate | Medium, High | Moderate |
| *Adelges tsugae* | Hemlock woolly adelgid | 41 | Herbivory, Chemical/physical/structural ecosystem impact, Interaction with other alien species, Other | Minor, Moderate, Major | Low, Medium, High | Major |
| *Apis mellifera* | European honeybee | 27 | Competition, Interaction with other alien species, Poisoning/toxicity, Other | Minimal Concern, Minor, Moderate, Major | Low, Medium, High | Major |
| *Bombus terrestris* | Buff-tailed bumblebee | 22 | Competition, Interaction with other alien species, Hybridisation, Transmission of disease, Other | Minimal Concern, Minor, Moderate | Low, Medium, High | Moderate |
| *Linepithema humile* | Argentine ant | 25 | Competition, Chemical/physical/structural ecosystem impact, Predation, Poisoning/toxicity, Facilitation of native species, Other | Minimal Concern, Minor, Moderate, Major | Low, Medium, High | Major |
| *Solenopsis invicta* | Red imported fire ant | 50 | Competition, Predation, Facilitation of native species, Interaction with other alien species | Minimal Concern, Minor, Moderate, Major | Low, Medium, High | Major |
